# High-resolution mapping of cell cycle dynamics during T-cell development and regeneration *in vivo*

**DOI:** 10.1101/2023.06.14.544919

**Authors:** Heike Kunze-Schumacher, Nikita A. Verheyden, Zoe Grewers, Michael Meyer-Hermann, Victor Greiff, Philippe A. Robert, Andreas Krueger

**Author notes:** Correspondence: A. K.; P. A. R. shared senior authorship (Philippe A. Robert, Andreas Krueger).

## Abstract

Control of cell proliferation is critical for the lymphocyte life cycle. However, little is known on how stage-specific alterations in cell cycle behavior drive proliferation dynamics during T-cell development. Here, we employed *in vivo* dual-nucleoside pulse labeling combined with determination of DNA replication over time as well as fluorescent ubiquitination-based cell cycle indicator mice to establish a quantitative high-resolution map of cell cycle kinetics of thymocytes. We developed an agent-based mathematical model of T-cell developmental dynamics. To generate the capacity for proliferative bursts, cell cycle acceleration followed a ‘stretch model’, characterized by simultaneous and proportional contraction of both G1 and S phase. Analysis of cell cycle phase dynamics during regeneration showed tailored adjustments of cell cycle phase dynamics. Taken together, our results highlight intrathymic cell cycle regulation as an adjustable system to maintain physiologic tissue homeostasis and foster our understanding of dysregulation of the T-cell developmental program.

## Introduction

Control of cell division is pivotal for development and function of the adaptive immune system. Effective B-cell responses in the germinal center as well as cytotoxic T-cell responses rely on rapid and massive clonal expansion of activated antigen-specific clones, which is mediated through alterations of the cell cycle^1–3^. However, substantially less is known about cell cycle control during developmentally programmed proliferation of lymphocytes.

The thymus produces naïve T cells with a diverse T-cell receptor (TCR) to fuel the adaptive immune system of jawed vertebrates over extended periods of life. In order to produce and select such a repertoire a large number of selectable precursor cells are generated during the course of T-cell development. Accordingly, it is characterized by a defined sequence of differentiation events interspersed by proliferative bursts. At steady-state the thymus is periodically colonized by bone-marrow (BM) derived thymus seeding progenitors (TSPs) filling a small number of niches in a CC chemokine-receptor (CCR)7 and 9 dependent manner^4–6^. TSPs give rise to early T lineage progenitors (ETPs), the most immature among CD4 and CD8 double-negative (DN) thymocytes^7,8^. ETPs progressively lose multi-lineage potential until T-lineage commitment is completed at the DN2b stage^9–11^. Subsequently, thymocytes rearrange their *Trb*, *Trg*, *Trd* loci to produce pre-TCRs or γδTCRs. Thymocytes expressing a pre-TCR enter the αβT-cell lineage marked by β-selection and transition from the DN3a to the DN3b stage. Following upregulation of the CD4 and CD8 co-receptors, double-positive (DP) thymocytes rearrange their *Tra* loci and undergo positive and negative selection to eliminate clones with signaling incompetent or potentially autoreactive T-cell receptors.

Studies on the turnover of thymocytes *in vivo* have determined the average residence time of thymocytes within phenotypically distinct populations^12–14^ and provided estimates of the number of cell divisions within populations^15,16^. After colonization, approximately 160 TSPs give rise to a population of 20,000 ETPs and 25,000 DN2 cells over a period of up to 14 days^4,13,17^. DN2 cells rapidly expand before proliferation ceases in DN3a cells to allow for TCR gene rearrangement. Proliferation then resumes after β-selection. At this stage, population size increases several hundred-fold to generate a sufficiently large pool of precursors for Tra rearrangement and subsequent selection events, which are accompanied by substantial amounts of cell death^18^.

The exact nature of how thymocytes adjust cell cycle length in a developmentally controlled manner or as a consequence of instructive signals, such as pre-TCR signaling remains steady-state and upon perturbation, we sought to generate a high-resolution map of cell cycle phase duration *in vivo*. To this end, we employed cell cycle indicator transgenic mice as well as dual-nucleoside pulse labeling combined with determination of DNA replication over time. The latter approach allowed us to discriminate cells almost immediately after entry into S phase as well as just prior to exit of S phase generating two virtually synchronized subsets within each thymocyte population. Thus, we determined exact cell cycle lengths at the level of individual cell cycle phases revealing proliferative heterogeneity directly *in vivo*. High-resolution data helped to establish a broadly applicable agent-based mathematical model (ABM) for cell cycle analysis, taking into account heterogeneous cell phases as well as *bona-fide* non-cycling cells. The model allowed us to extract the duration of all phases while consistently explaining the dynamics of all labeled populations, and assign heterogeneous behavior to distinct cell cycle phases. Finally, we applied our method to a model of thymus regeneration and showed that different thymocyte populations modulate both S-phase and G1-phase duration with distinct patterns.

## Results

### Single-nucleoside pulse labeling and cell cycle reporter mice reveal composition and heterogeneity of cell cycle stages in thymocyte populations

We employed BrdU single-pulse labeling combined with analysis of DNA content to determine the steady-state proportions of wild-type (WT) thymocytes at distinct cell cycle states (Fig. 1a,b). In all subpopulations the majority of cells were in the G0/G1 phase of the cell cycle (BrdU^-^, 2N DNA) and a small fraction of cells was in G2/M phases (BrdU^-^, 4N DNA) (Fig. 1c). The largest proportion of DNA replicating cells (S phase, incorporation of BrdU and 2N < DNA content < 4N) were found in the DN2, DN3b and DN4 subsets (Fig. 1c), whereas the lowest proportions were found in DN3a and pre-selection DP cells as well as post-selection DP and SP thymocytes (Fig. 1c).

**Figure 1.**
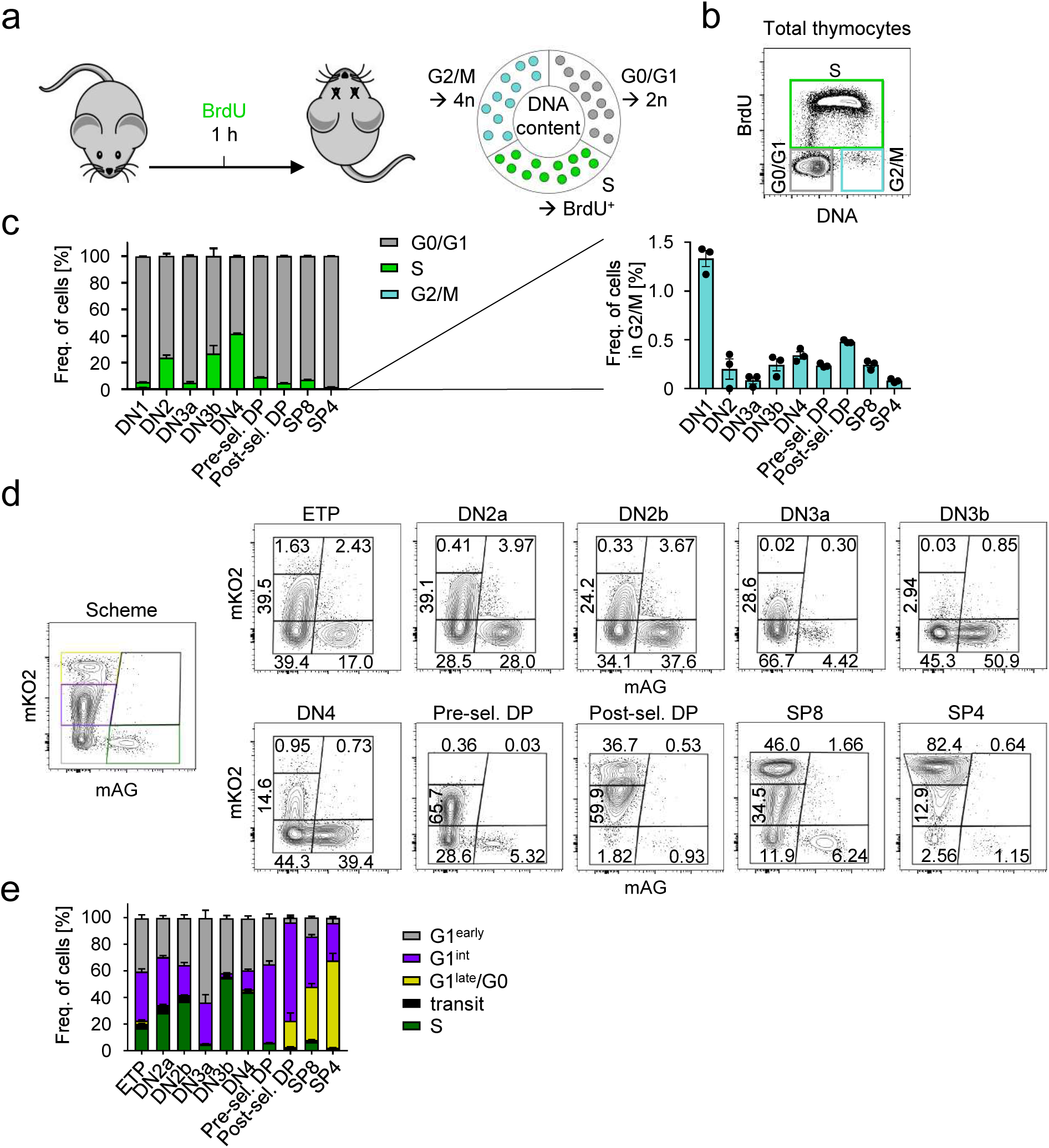
Single-nucleoside pulse labeling and cell cycle-reporter mice reveal composition and heterogeneity of cell cycle-stages in thymocyte populations. **(a)** Schematic representation of the performed experiments using *in vivo* BrdU single-pulse labeling in combination with analysis of DNA content. **(b)** Representative flow cytometric gating strategy to identify cells in G0/G1-, S- or G2/M-phase. **(c)** Frequencies of cells in G0/G1-, S- or G2/M-phase of murine WT thymocyte subpopulations, n = 3 mice. **(d)** FUCCI gating strategy and profiles of WT thymocyte subpopulations. Gates were defined as G1^early^ (grey, mAG^-^mKO2^-^), G1^int^ (purple, mAG^-^mKO2^+^), G1^late^/G0 (yellow, mAG^-^mKO2^++^), transit (black, mAG^+^mKO2^+^) and S (green, mAG^+^mKO2^-^). **(e)** Frequencies of FUCCI cell subsets as defined in (d) of WT subpopulations, n = 6 mice.

Next, we analyzed a transgenic mouse model with fluorescent reporters linked to cell cycle progression (FUCCI mice)^19^. These mice express a green fluorescent reporter (mAG) during S phase, whose frequencies in thymocytes correlated well with S-phase frequencies determined by single BrdU pulse and DNA content analysis, indicating faithful reporting of cell cycle stages in the FUCCI system (Fig. 1d,e). Upon transition through G2/M phases back to G1 phase, cells rapidly lose mAG expression through proteolytic degradation followed by a continual increase in expression of an orange fluorescent reporter (mKO). High levels of mKO are therefore indicative of an extended G1 phase or even quiescence^19^. By assuming that mKO expression increases at the same speed in all populations, the fraction of cells with low or intermediate mKO levels (early in the G1 phase) provide qualitative information on G1-phase duration and heterogeneity. We subdivided mAG-negative thymocyte populations according to mKO levels into three subsets, mKO-negative/low, mKO-intermediate, and mKO-high, representing cells in G2/M/early G1 (G1^early^), intermediate time spent in G1 (G1^int^), and late G1/G0, respectively. ETPs and downstream populations up to DN3a thymocytes contained between 20 to 40 % of cells in G1^int^, while DN3b and DN4 showed lower amounts of G1^int^ cells, 3% and 15% respectively, indicating that DN3b and DN4 exhibit shorter G1 phases compared to the former populations. Thus, FUCCI reporter expression suggests that rapidly cycling DN3b and DN4 thymocytes are characterized by a higher frequency of cells in S phase as well as reduced time spent in G1 phase (Fig 1d,e). Post-selection thymocytes progressively acquired high and discrete levels of mKO, suggesting that these cells ultimately exit from the cell cycle on the path of maturation towards naïve T cells (Fig. 1d,e). Taken together, steady-state analysis confirmed previously established switches between non-proliferative and proliferative phases of T-cell development. Different expression levels of the G1 reporter in FUCCI cells suggested that similarly replicating populations, such as DN2 and DN3b cells, are nevertheless likely to exhibit distinctive cell cycle behavior.

### Dual-nucleoside pulse labeling reveals rates of entry in S phase

Next, we employed dual-nucleoside pulse labeling to study dynamic cell cycle behavior. Cells were sequentially labeled *in vivo* with EdU and BrdU at 1 h apart followed by analysis 1 h after the second pulse (Fig. 2a). The window of action of EdU or BrdU labeling over detection threshold has been estimated to be 45-60 min after label administration, with EdU displaying a somewhat shorter bioavailability than BrdU^20^. Thus, labeling at a 1 h interval creates single-nucleoside positive EdU^+^BrdU^-^ and EdU^-^BrdU^+^ populations, representing cells that have ceased DNA replication within 1 h after label administration (post S) and cells having initiated DNA replication within less than 1 h after BrdU administration (early S), respectively (Fig. 2b). Thus, single-nucleoside positive populations provide information on rates of entry into and exit from S phase. Cells without label (EdU^-^BrdU^-^) did not go through DNA replication during the period of EdU and BrdU labeling, and were either in G1, G2/M or potentially quiescent.

**Figure 2.**
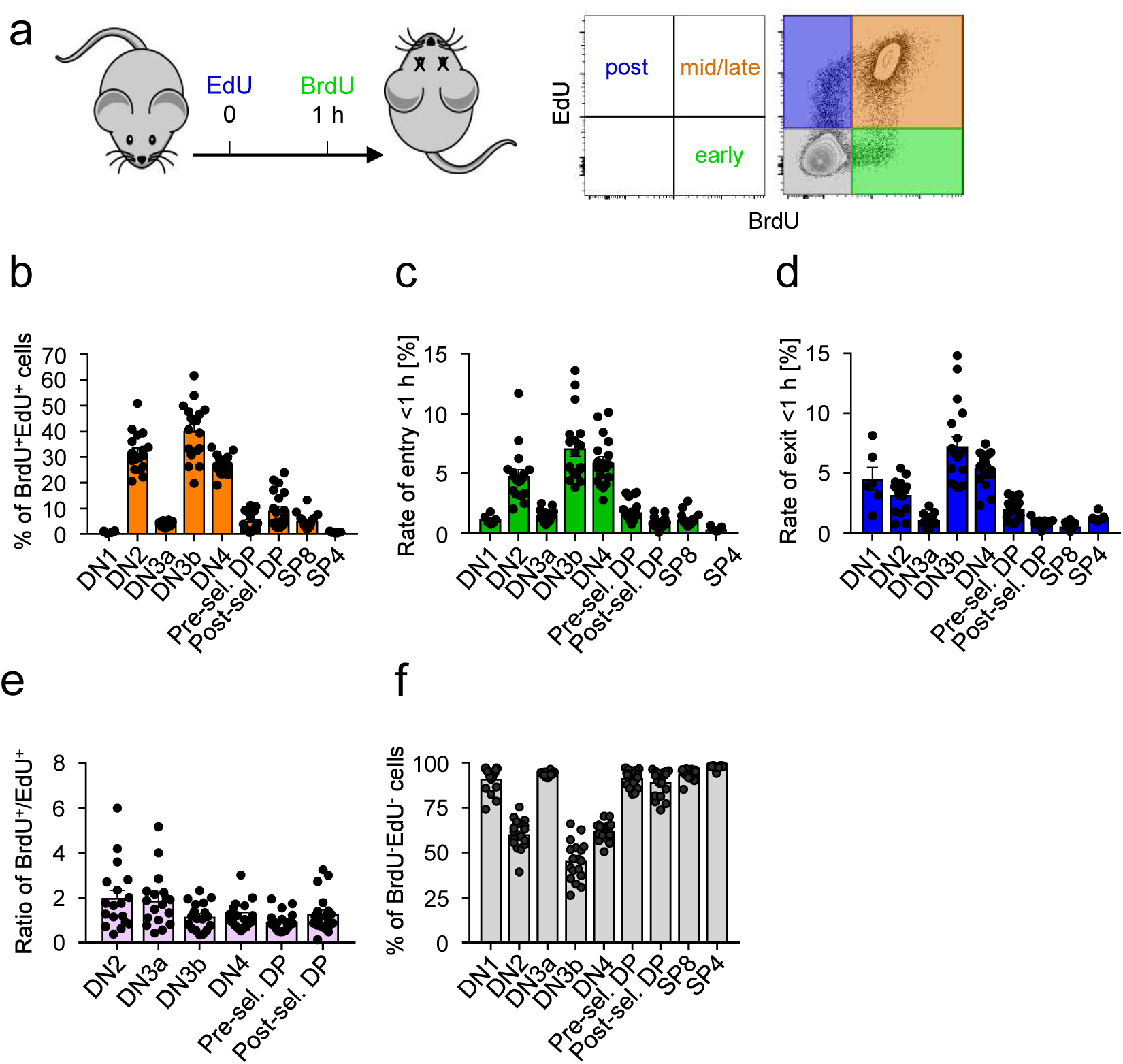
Dual-nucleoside pulse labeling reveals rates of S-phase entry. **(a)** Schematic overview and representative flow cytometric gating strategy of *in vivo* dual-pulse labeling. Mice received consecutive pulses of EdU and BrdU 1 h apart. Analyses were performed 1 h after the second pulse. Thymocytes in S phase during both pulses appear as double positive, EdU^+^BrdU^+^ (orange, mid/late S phase). EdU- or BrdU-single positive thymocytes represent cells which have ceased DNA replication and left S phase (EdU^+^, blue, post S phase) or newly entered S phase during the second pulse (BrdU^+^, green, early S phase), respectively. **(b-e)** Graphs show frequencies of EdU^+^BrdU^+^ cells **(b)**, EdU^-^BrdU^+^ cells **(c)**, EdU^+^BrdU^-^ cells **(d)** and EdU^-^BrdU^-^ cells **(e)** of murine WT thymocyte subpopulations, n = 6-20 mice of two independent experiments.

Analysis of EdU^-^BrdU^+^ thymocytes showed that DN2, DN3b, and DN4 had frequencies of S-phase entry of 5-7% in <1 h (Fig. 2c). DN1, DN3a, pre-selection DP cells had frequencies of S-phase entry around 2% in <1 h and those of post-selection DP and CD8SP cells were even lower (Fig. 2c). Virtually no CD4SP cells started DNA replication within this time frame (Fig. 2f). Taken together, the frequency of thymocytes starting DNA replication corresponds well with the steady-state analysis (Fig. 1c) in that cells with higher frequency of DNA replication also show higher hourly frequencies of S-phase entry.

Frequencies of EdU^+^BrdU^-^ cells, corresponding to cells exiting S phase during the labeling process, were comparable to those of EdU^-^BrdU^+^ thymocytes, but somewhat higher throughout (Fig. 2d, e), which might be due to small differences in bioavailability of both nucleoside analogs. In theory, the amount of cells entering and exiting the S phase should be equal, except if cells differentiate, die or exit within the S phase. DN1 and CD4SP cells displayed considerably higher exit than entry rates. However, due to the low cell numbers within the EdU^+^ subsets, definitive conclusions for possible underlying mechanisms are difficult to draw. One may speculate that in the case of CD4SP cells, EdU^+^ cells represent the final cycle before cells enter final maturation and the frequency of quiescent cells increases. Additionally, it is possible that in these populations DNA replication follows different kinetics early or late during S phase, resulting in divergent levels of nucleoside incorporation.

Assuming that in all thymocyte populations progression through the cell cycle is homogeneous, i. e., times between each round of DNA replication are identical within a thymocyte population, rates of S-phase entry can be converted into cell cycle duration: the percent of cells entering the S phase in 1 h informs how many hours are necessary for 100% of the initial cycling cells to have entered the S phase and therefore provides an estimate of the full cycle (Fig. 2c). Due to the virtual lack of quiescent cells identified in G0 in all subsets except SP thymocytes (Fig. 1d), it can be first assumed that all cells within one subset are cycling, although in part a substantial fraction of cells does not acquire any label with two sequential pulses. Under this assumption, DN2, DN3b and DN4 cells would complete one full cycle within 12 h, 8 h, and 9.5 h, respectively, and thus represent the subsets with the shortest cycling periods. Longer cell cycle periods were estimated for DN1, DN3a, as well as pre-selection DP with average cell cycle durations of 47, 36, 34 h, respectively. Post-selection DP and CD4SP thymocytes had estimated cell cycle durations that extended well beyond their estimated life times, while CD8SP were estimated to cycle in 52 h, consistent with their progression to quiescence and showing that the rate of S-phase entry cannot directly be translated into cell cycle duration due to presence of quiescent cells^17^.

Taken together, steady-state analysis of cell cycle using single-pulse and dual-pulse nucleoside labeling revealed substantial differences between thymocyte populations with slow and rapid turnover, consistent with previous studies^15^. They also provided initial quantitative information about total cell cycle duration based on the determination of the rate of cells starting DNA replication. Furthermore, dual-pulse labeling provided qualitative information on the relative contribution of the S phase to the overall cell cycle. Nevertheless, it remains unclear to what extent cells perform the cell cycle in a homogeneous and synchronized manner and therefore whether direct estimation of cell cycle from steady-state and rates of entry or exit is possible.

### High resolution tracking of virtually synchronized cells using dual-pulse labeling combined with DNA content analysis over time informs on the duration of individual cycle phases

Next, we sought to quantitatively determine the duration of cell cycle stages in each thymocyte subset. Dual-nucleoside pulse labeling at 1 h apart as described above creates virtually synchronized populations at the start of S phase (EdU^-^BrdU^+^) and at its end (EdU^+^BrdU^-^). DNA content analysis was performed to trace S-phase progression of virtually synchronized cells from 1 hour to 20 hours post BrdU injection (Fig. 3a). Due to limitations in cell numbers and few nucleoside incorporating cells, DN1 and both SP subsets were omitted from this analysis.

**Figure 3.**
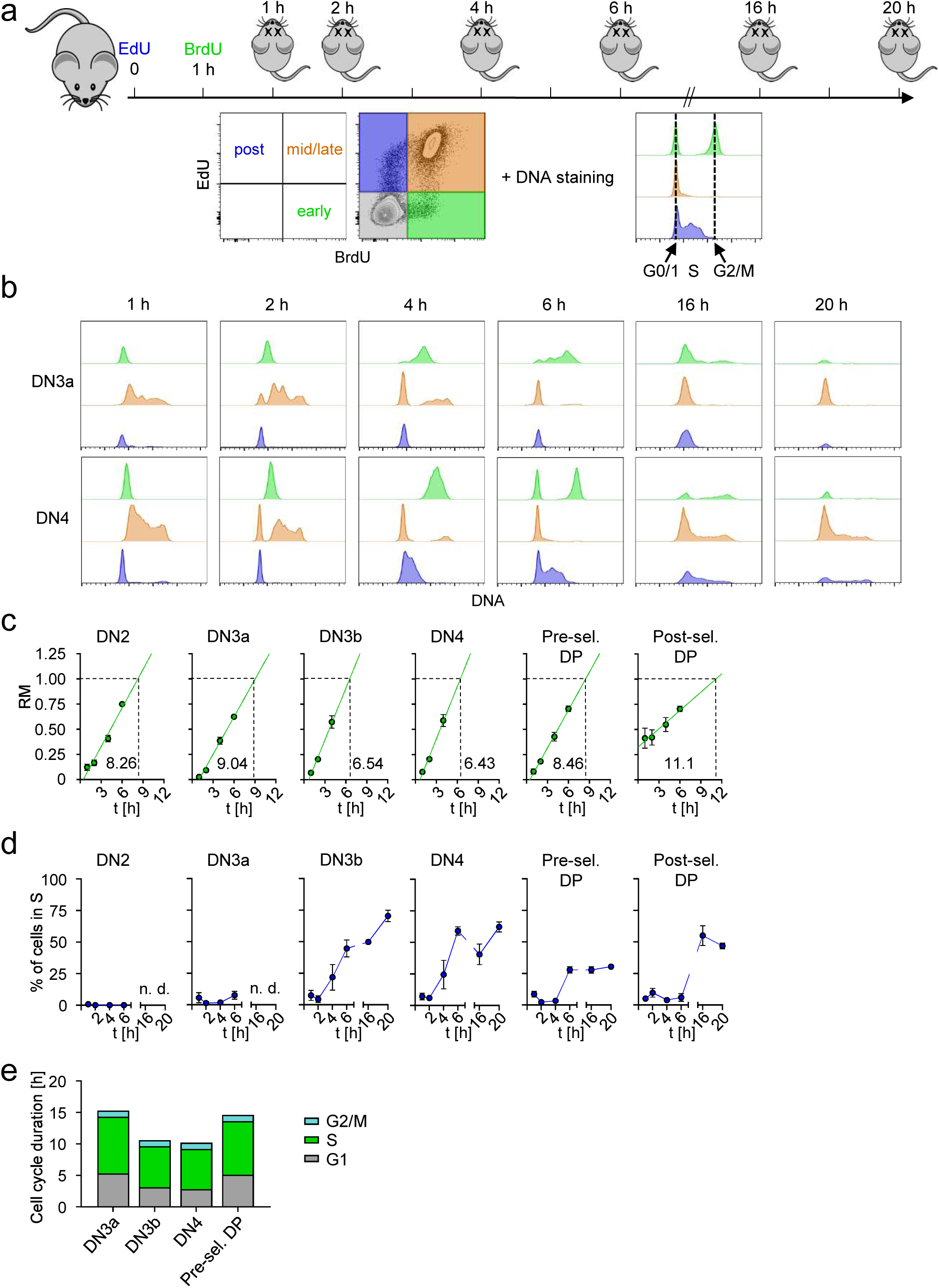
High resolution tracking of virtually synchronized cells using dual-pulse labeling combined with DNA content analysis over time informs on the duration of individual cycle phases. **(a)** Schematic overview and representative flow cytometric gating strategy of *in vivo* dual-pulse labeling. Mice received consecutive pulses of EdU and BrdU 1 h apart as described in Figure 2. Subsequent analyses were performed 1, 2, 4, 6, 16 or 20 h after the second pulse. DNA staining was performed to identify cell cycle-states. **(b)** Representative flow cytometric histograms visualize the DNA content of DN3a and DN4 thymocytes of WT mice over time. Each plot depicts an overlay of the DNA content of EdU^-^BrdU^+^ (green), EdU^+^BrdU^+^ (orange) and EdU^+^BrdU^-^ (blue) cells. **(c)** Statistical analysis of WT thymocyte subpopulations to assess S-phase duration based on RM values of EdU^-^BrdU^+^ cells (early S phase) over time (green dots). The green line represents a linear regression. Numbers adjacent to linear regression show S-phase time in h calculated based on the linear regression, n = 3-5 mice for each time point, data from 2 independent experiments. **(d)** Quantification of S-phase re-entry of EdU^+^BrdU^-^ WT thymocyte subpopulations over time, n = 4-5 mice for each time point, data from 2 independent experiments. **(e)** Total cell cycle-length and cell cycle-phase composition of different WT thymocyte subpopulations. For DN3a, DN3b, DN4 and pre-selection DP thymocytes, length of G1 phase was calculated based on RM values of EdU^+^BrdU^-^ S-phase cells at later time points (dark grey). S-phase length (green) was determined as shown in (c).

The dynamics of DNA incorporation of cells that started (green), exited (blue) or stayed (orange) in S phase during the interval of the first labeling is shown for a representative slow- and fast-proliferating population, DN3a and DN4, respectively (Fig. 3b, Fig. S1a).

We determined S-phase duration in virtually synchronized cells at S-phase entry (EdU^-^BrdU^+^, green) by monitoring the increase of DNA content from 2N to 4N towards a complete round of DNA duplication (Fig. 2c). A similar analysis with dual-labeled cells (EdU^+^BrdU^+^, orange), which are non-synchronized and hence cover a broad range of S-phase states, yielded similar results (Fig. S1b)^21^. We determined S phases of 6-7 h for DN3b and DN4 cells, consistent with rapid turnover of these populations (Fig. 3c). Longer S-phase duration of 8-9.5 h was observed for DN2, DN3a and pre-selection DP cells, although DN2 cells displayed higher rates of cell cycle entry compared to DN3a and DP, and were therefore cycling substantially faster than the latter two populations (Fig. 2c). These data suggest that S phase shortening below a certain limit is a feature restricted to extremely fast proliferating DN3b and DN4 subsets. Curiously, post-selection DP cells contained more DNA already at early time points of analysis and cells remained in S phase for considerably longer than other populations. The underlying mechanism of this peculiarity remains unknown.

Virtually synchronized cells at S-phase exit (EdU^+^BrdU^-^) had mostly reached G1 already at the 1 h time point of analysis, corresponding to 1.25 – 1.5 h after exiting S phase, if we include a delay of labeling bioavailability (Fig. 3b, Fig. S1a, blue histograms). We conclude that for all analyzed thymocyte populations G2/M duration is less than 1.5 h. Pre-selection and post-selection DP cells had somewhat higher frequencies of cells remaining in G2/M at this point in time, suggesting that the distribution of G2/M may be broader or that the transition between S and G2/M phases is prolonged (Fig. S1a). G1 phase duration can be derived from determining the time point of the onset of DNA replication (DNA content >2N) of cells virtually synchronized at S-phase exit (Fig. 3b, Fig. S1a, blue histograms). DN3b and DN4 populations displayed a minimal G1-phase duration of 3.1 and 2.8 h, respectively (Fig. 3d). DN3a and pre-selection DP cells had minimal G1 phase durations of 5.25 and 5.1 h (Fig. 3d). Surprisingly, G1 phase duration in DN2 cells extended beyond this type of analysis. DNA content analysis after 16 and 20 h revealed plateaus of DNA content at approximately 50% for DN4 and post-selection DP cells and 30% for pre-selection DP cells. This analysis assumes identical cell phases durations. If the G1 duration is heterogeneous between cells, this approach may focus on cells with the shortest G1 duration and underestimate the population average G1 duration. Apparent stable frequencies of G1 cells at later time points can be explained by de-synchronization of cell cycles over extended periods of time, transitions to the subsequent developmental stage or both.

We tested whether label-negative cells provided some indication of population heterogeneity. Consistent with estimated cell cycle durations, cell cycle entry in at least some label-negative cells was detected already at 2 h in fast cycling populations (DN3b and DN4) and at 4 h in DN2 cells. In contrast, no cell cycle entry was observed in pre-selection DP cells, consistent with these cells ceasing proliferation in order to start TCR gene rearrangements (Fig. S2).

Taken together, combination of dual-pulse labeling with analysis of temporally resolved DNA replication provided a direct estimate on the duration of individual cell cycle phases for thymocytes (Fig. 3e). Partially overlapping information obtained from different experiments allowed us to more precisely define boundaries of cell cycle duration for most subsets. Comparing the different approaches of cell cycle mapping, we identified cell populations with highly congruent quantitative data across setups, such as DN3b and DN4 thymocytes. For other populations, cell cycle phase level data generally indicated shorter cycle durations, when compared to inferred cell cycle durations from steady-state data. These apparent discrepancies point towards intra-population heterogeneity, which was further explored through mathematical modeling and experimental validation.

### Modeling thymocyte population dynamics based on high-resolution cell cycle analysis

To integrate apparent quantitative discrepancies between the analyses proposed above, and to accurately estimate the duration and heterogeneity of the cell cycle phases from dual-labeling experiments combined with DNA content analysis, we developed an ABM of cell division and labeling. The model simulates a population of cells progressing through the G1, S and a combined G2/M phase that includes both the G2 phase and mitosis (Fig. 4a, left). Inspired by the Cyton model^22^, the duration of each phase follows a lognormal time distribution whose average and width are to be estimated from experimental data (the unknown parameters). When a cell divides, two new daughters are created at G1, and a time is picked from these distributions for the ending of G1, S and G2/M phases. A death time is also sampled at birth from an exponential time distribution, and death is triggered if it happens before the end of G2/M. We incorporate a separate pool of cells whose cycle is stopped for a long time (either in G0 or G1), that we call ‘long G1’ cells.

**Figure 4.**
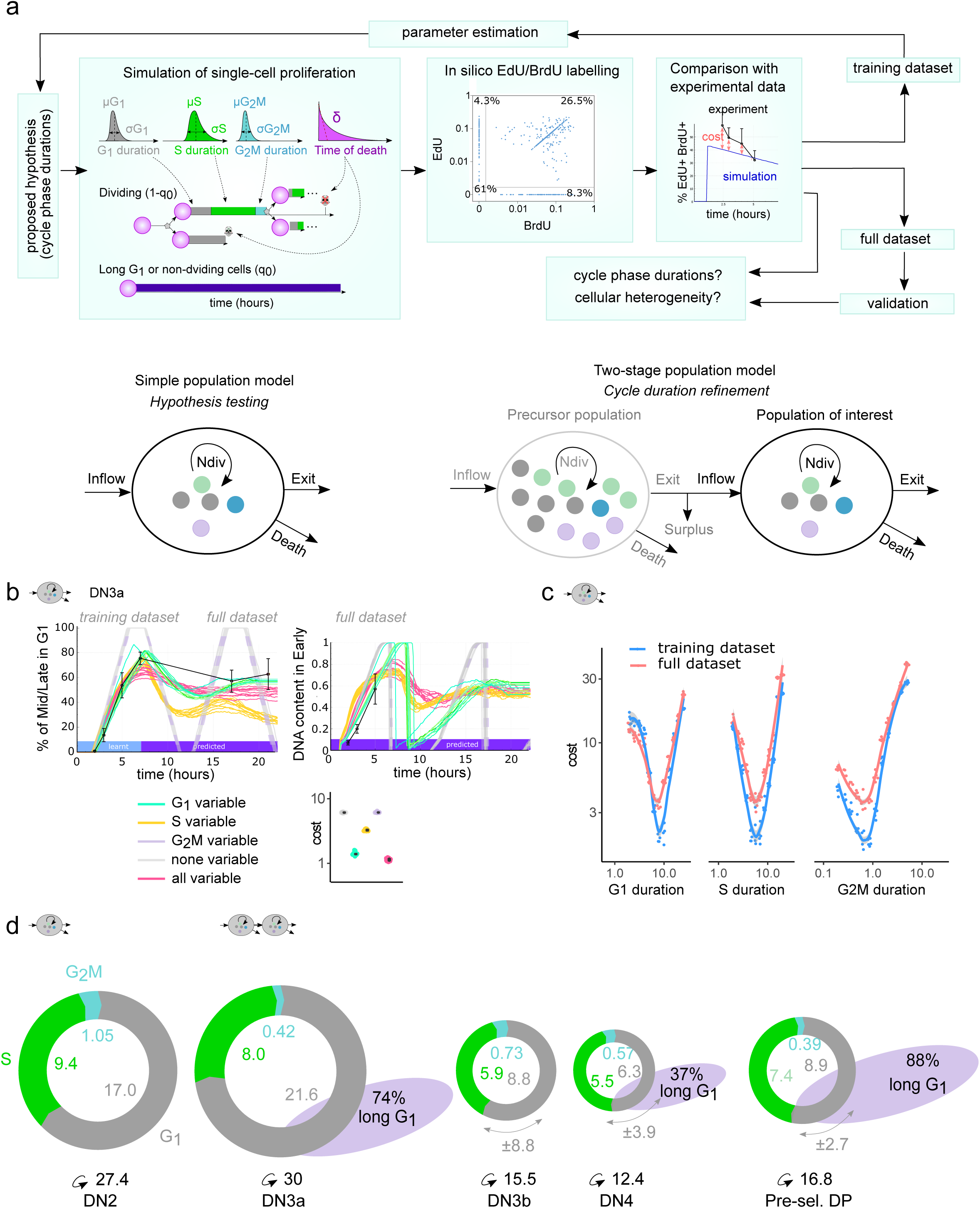
Modeling thymocyte population dynamics based on high-resolution cell cycle-analysis. **(a)** Workflow for phase duration inference. Left: The agent-based mathematical model (ABM) simulates cell cycle-progression using lognormal time distributions for phase durations and a constant death rate as an exponential distribution. A fraction of bystander cells that do not incorporate EdU nor BrdU, termed ‘long G1 cells’ can be considered. Center: *In silico* single-cell profiles for EdU and BrdU labeling mirror those generated experimentally by flow cytometry. Right: A cost (likelihood) evaluates the quality of a simulation compared to the data. Parameter estimation algorithms iteratively find the cell cycle-durations that best explain the data. Bottom: The model simulating one population is used for hypothesis comparison, while the two-populations model is used for a refined estimation of cycling speed when the phase durations of the ancestor populations are known. The two-populations model is used to validate that inferred cell cycle phase durations are not sensitive to incorporating additional levels of biological complexity such as inflow of already labeled cells from the precursor population. **(b)** Comparison of cell cycle-heterogeneity hypotheses for the DN3b population. ‘none variable’ hypothesis: all cells share the same duration for a phase. Under other hypotheses, one or multiple phases are allowed to be variable between cells. The cost and curves of 10 independent fittings for each hypothesis are shown (see Figure S3_1 and S3_2 for breakdown of curves under different hypotheses). Training data-points are annotated in light blue (‘learnt’), and those only in the validation dataset are annotated in dark blue (‘predicted’). For the DN3b population, a heterogeneity in the G1 (but not other phases) is required to explain the curves with minimal cost. The hypothesis ‘all variable’ does not improve the quality of the simulation compared to the data. **(c)** Identifiability analysis of the found phase durations by profile likelihood for the DN3b population: the cost of the simulations is shown after fixing the value of each cycle phase duration separately (x axis) while other cycle phases are estimated. A minimum in a curve shows that there is a unique cell cycle duration that best explains the data, meaning that the dataset was rich enough to identify the phase duration. A flat curve would mean that different values for the phase duration (x axis) explain the data equally well (same cost), i.e. the duration cannot be identified. Blue: training dataset only; Red: train and test datasets. The cell cycle durations (curve minima) found using the training dataset are the same as those found with the full dataset. **(d)** Cell cycle phase durations identified using the full dataset, with populations. The identified percent of ‘long G1’ cells is shown in gray and the estimated variation of G1 phase durations are shown with the double arrow.

We reproduce the dual pulse EdU and BrdU labeling as in the experimental settings (Fig. 4a, center). Cells are gated *in silico* as in flow cytometry into four populations: unstained (EdU^-^BrdU^-^), EdU^+^BrdU^-^, EdU^-^BrdU^+^ and EdU^+^BrdU^+^ populations. Cells from each gate are assessed for their cell cycle phase. A cost is calculated between a simulation and the experimental data (Fig. 4a, right, Methods). Automated parameter estimation iteratively simulates labeling for many possible hypothetical durations of each cell phase, and identifies the most likely cell cycle durations by minimizing the cost. For each population, we separated the experimental data in a training dataset containing the first four analysis time points for parameter estimation for all variables except DNA levels, and kept DNA levels and the remaining time points for validation. First, we simulated one population at a time to compare different hypotheses of cell cycle heterogeneity. Second, we expanded this model by adding simulations of the labeling dynamics of cells from the immediate precursor population, thus creating a two-populations model (Fig. 4a, bottom). We employed the latter model to provide a refined estimate of cell cycle phase durations.

To avoid overfitting due to unnecessary model complexity, we devised five different hypotheses based on heterogeneity of the cell cycle phases between cells of the same population (Fig. 4b) and performed parameter estimation 10 independent times on the training dataset. Each parameter estimation takes between 4 h and 6 h on a single laptop CPU in the one-population model. The simplest hypothesis (‘non-variable’), assumes a fixed duration of each phase. Intermediate hypotheses (‘G1 or S or G2/M variable’) consider only one phase to be variable between cells, while the most complex (‘all variable’) allows each phase to have its own width. For each population, we compared the quality (cost) of each strategy, as well as the presence of bystander ‘long G1’ cells. In the example of the DN3b population, the hypotheses ‘variable G1’ and ‘all variable’ showed a lower cost than the other hypotheses (Fig. 4b, Fig. S3_1, Fig. S3_2). Interestingly, the hypothesis ‘variable S’ did not perform better than ‘variable G1’, and was inconsistent with the “%Post in G1’’ and the DNA labeling dynamics, suggesting that variable S phase durations are not required to explain the data. The ‘all variable’ hypothesis did not further improve the quality of the curves nor the cost. Thus, we concluded that a variable G1 phase provides the best explanation for the data with minimal model complexity in DN3b cells. Of note, this finding was confirmed in the full validation dataset (Fig. S3_3). The simulation curves with parameters inferred from the training dataset using the ‘variable G1’ hypothesis were overall largely consistent with the labeling dynamics at the later time-points (Fig. S3_2). For comparison, the curves obtained when inferring parameters from the full dataset with the ‘variable G1’ were not substantially better than the ones from the training dataset, indicating that the later time-points may contain certain levels of biological complexity that the model can not perfectly explain (Fig. S3_2).

We confirmed that cell cycle phase durations can confidently be identified using a profile likelihood analysis, which compares the cost of parameter estimations after fixing a phase duration to different possible values on the training dataset (Fig. 4c, Fig. S3_6, blue curves). The existence of a minimum cost means that only one value for this phase duration best dataset. Alternatively, a flat curve would mean that different durations would explain the dataset equally well and are therefore nonidentifiable. We conclude that the duration of all three phases is accurately identified by our approach. For the DN3b (Fig. 4c), DN4 and pre-sel DPs (Fig. S3_6), four time-points (training dataset, blue curves) were sufficient to identify cycle phase durations, and the identified minima were the same in the training and the full dataset (red curves). However, for the DN2 and DN3a populations, a minimum of 6 time-points was required to identify G1 duration because those populations have a longer G1 phase that cannot be captured by the initial four time-points. We also confirmed that the settings of other simulation parameters than phase duration did not significantly impact the results by performing an identifiability analysis varying the death rate or the number of divisions (Fig. S3_4). Therefore, the combined experimental-modeling approach is able to identify phase heterogeneity and duration *in vivo,* and we provide the cell cycle phase durations for each population based on the best-suited hypothesis for each population (Table S1).

These simulations were based on a homogeneous one-population model seeded by fresh unlabeled cells, thereby ignoring that entering cells from the precursor population may already be labeled by EdU or BrdU. To assess the effect of inflow of such labeled cells, we developed a two-stage model simulating dual-pulse labeling for both the population of interest and its precursor population, and in which newcomer cells come from cells exiting the precursor population (Fig. 4b). This model provides the flexibility of deciding whether precursors enter the main population after mitosis or at a random phase of the cell cycle, but is extremely slow since we need to simulate a large population of precursors to be certain to provide sufficient inflow to the next population (five days for one parameter estimation on a single CPU), which makes identifiability analysis and hypothesis comparison impossible. The phase durations of a population are optimized while fixing the precursor phase durations from those identified by the one-population model. The estimated cell cycle durations were similar between both the one-population and the two-populations models (Fig. S3_6), demonstrating that the complexity of the two-populations model is not required per se to identify cell cycle durations, but can be used as a refinement that incorporates more biological complexity.

We show the refined cell cycle phase durations identified by the two-populations model based on the best-suited hypothesis for each population (Fig. 4d, Table S1). The DN2 and DN3a labeling kinetics could be explained without phase variation, while the other populations benefited from variation of the G1 phase (Fig. S3_3). The cycle durations ranged between 12.4 h for DN4 to 30 h in the DN3a population. The S phases ranged between 5.5 h explained without ‘long G1’ cells, which is expected for these populations, while other populations needed a substantial amount of such cells. The DN3a population required 74% of ‘long G1’ cells, consistent with the need for TCR rearrangement and β-selection at this stage. The pre-selection DP population was best explained including 88% of ‘long G1’ cells, consistent with proliferation arrest TCRα gene rearrangement and onset of thymic selection at this stage. Surprisingly, the DN4 population required ∼37% of ‘long G1 cells’, which was necessary to explain the low re-entry of unlabeled cells into the S phase, suggesting the existence of a subpopulation of cells with different cell cycle kinetics.

### Decision to re-enter S phase after one round of cycling defines cell cycle heterogeneity

To experimentally validate heterogeneous behavior of cells we first subjected sorted DN3a thymocytes at defined stages of the cell cycle to short-term *in vitro* differentiation on OP9-DL1 co-cultures (Fig. 5a). FUCCI negative (corresponding to early G1) cells progressively acquired mKO2 fluorescence over time, in line with prolonged time spent in G1 (Fig. 5b,c). A subset of cells (2 – 4%) in S phase (mAG positive) was detected already after 2 h of culture and the frequency of this population remained virtually constant throughout the period of culture, suggesting that this population is not continually fueled by cells from mid/late G1 phase. Sorted mKO2-positive cells (mid/late G1) gave rise to a larger proportion of S phase cells over time when compared to early G1 cells, increasing over a period of 12 h, but remaining constant thereafter (Fig. 5d). Early G1 cells appeared after 14 to 16 h of culture, consistent with the S-phase duration of 7.5 to 9 h determined *in vivo*, and reached a plateau after 20 h. Starting from sorted mAG-positive (S phase) cells, frequencies of early G1 cells steadily increased between 2 and 12 h, followed by an expansion of intermediate G1 cells, indicating that intermediate G1 cells emerge from FUCCI negative cells after approximately 10 h (Fig. 5e). In contrast, no *de novo* increase in S phase cells was observed during the culture period. Taken together, these data indicate that the transition from S phase through division into G1 is kinetically fixed, whereas re-entry into S phase is independent from a predetermined G1 phase duration.

**Figure 5.**
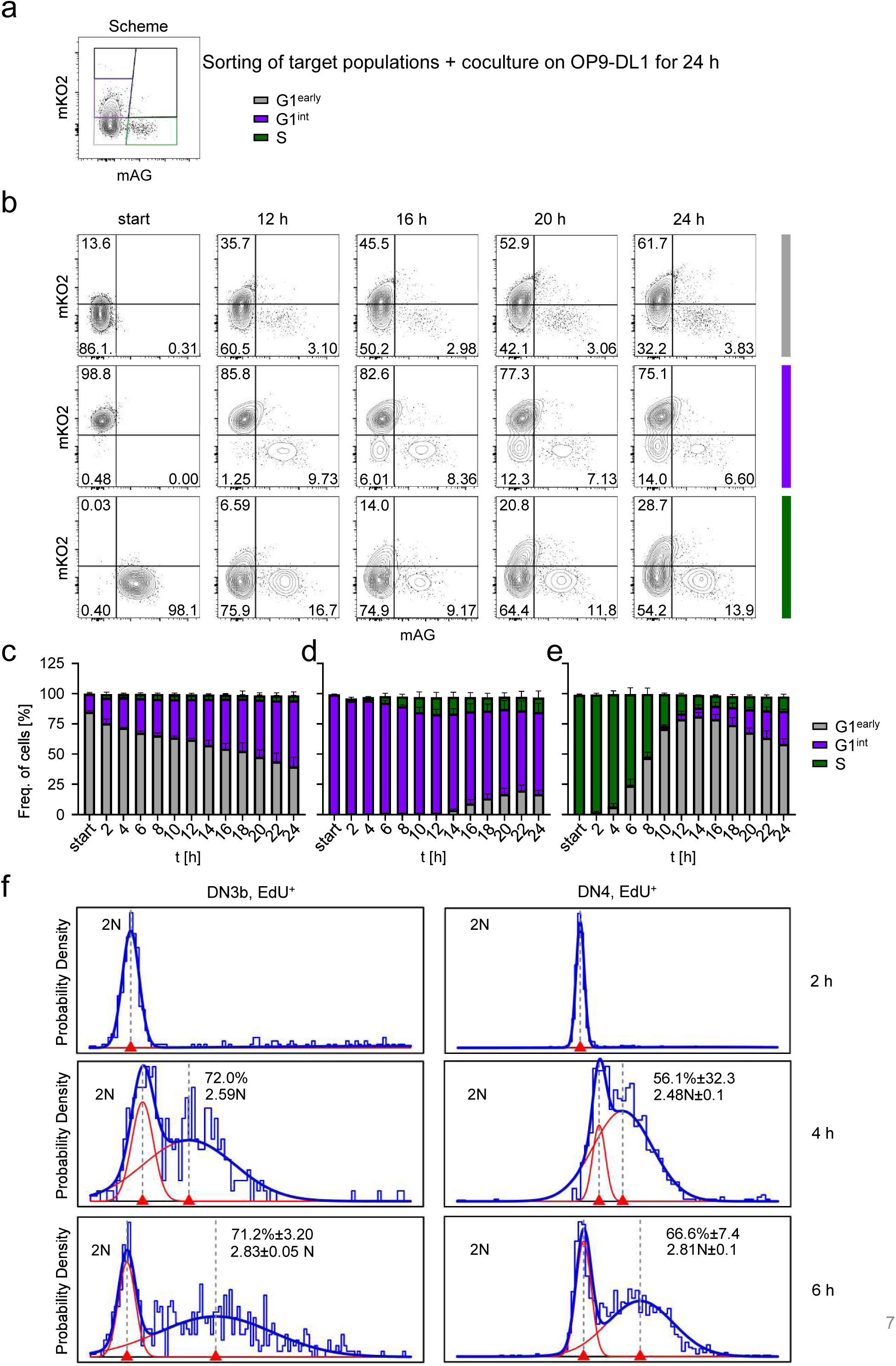
Decision to re-enter S phase after one round of cycling defines cell cycle heterogeneity. **(a)** Schematic representation of experiments. FUCCI DN3a thymocytes were sorted based on the presented gating strategy as G1^early^ (grey, mAG^-^mKO2^-^), G1^int^ (purple, mAG^-^mKO2^+^) and S (green, mAG^+^mKO2^-^). Target populations were cocultured on OP9-DL1 feeder cells for up to 24 h and FUCCI cell cycle profiles were assessed by flow cytometry every 2 h. **(b)** Representative dot plots of G1^early^ (grey, mAG^-^mKO2^-^), G1^int^ (purple, mAG^-^mKO2^+^) and S (green, mAG^+^mKO2^-^) DN3a thymocytes at indicated time points. **(c-e)** Graphs show frequencies of cells of the indicated target populations and their corresponding FUCCI profiles over time with **(c)** G1^early^ (grey, mAG^-^mKO2^-^), **(d)** G1^int^ (purple, mAG^-^mKO2^+^) and **(e)** S (green, mAG^+^mKO2^-^). Pooled data from two independent experiments. **(f)** Identification of speed of DNA incorporation in cells re-entering the S phase. The distribution of DNA in the populations of interest is shown in logarithmic scale and rescaled between 2N and 4N using 2 h as reference for 2N. A mixture of two gaussians of unknown mean and standard deviation were fitted to the distribution using the mixdist R package. The percentage of cells as well as the average DNA level of cells covered by each gaussian is shown for the displayed data (n = 1-3).

Next, we assessed virtually synchronized thymocytes from dual-pulse labeling experiments for evidence of intra-population heterogeneity. Experiments described above allowed us to faithfully quantitate the duration of one round of cycling, but did not provide information about multiple division cycles. Here, we analyzed the onset of a second round of DNA replication in virtually synchronized fast cycling DN3b and DN4 cells at S-phase exit (Fig. 5f). Both populations displayed a bimodal distribution of DNA amounts detectable at 4-5 h after completion of the previous S phase. Whereas approximately 70 and 60 % of DN3b and DN4 cells, respectively, had passed G1 and re-initiated DNA replication at that time point, the remainder of cells were in G1 phase. Of note, this bimodal distribution remained essentially constant for at least an additional 2 h.

Taken together, *in vitro* and *in vivo* data validated cell cycle heterogeneity predicted by our mathematical model and show that, at least for DN3b and DN4 cells, this heterogeneity predominantly manifests itself prior to or at the G1 to S phase transition.

### Population-specific cell cycle phase alterations determine thymus regeneration

Restoration of thymus size after pre-conditioning constitutes a major bottleneck for regeneration of the immune system after hematopoietic stem cell transplantation^23^. We determined cell cycle dynamics during regeneration of thymus size at 6 days after sublethal irradiation, when most populations have recovered to at least 50% of steady-state (Fig. 6a, Fig. S5_1a). First, we performed dual-pulse labeling to determine frequencies of cells in S phase as well as rates of S-phase entry as a measure of total cell cycle length. At 6 days post-irradiation, all populations with the exception of DN3a cells displayed substantial frequencies of cells in S phase, ranging from 27 – 69% (Fig. 6b). When compared to the corresponding populations at steady-state in non-manipulated mice, DN2 and DN3a cells had 1.5 – 2.5-fold higher frequencies in S phase, whereas these frequencies were unaltered in DN3b and DN4 cells (Fig. 6b). Most notably, both subsets of DP thymocytes displayed a 4.6 – 26-fold increase of S phase cells when compared to the steady state. Entry rates into S phase during regeneration were similar in DN2 and DN3b cells, when compared to steady-state, but markedly increased in DN3a, DN4 and pre-selection DP thymocytes, indicating shorter overall cell cycle duration in these populations (Fig. 6c). Whereas at steady-state S-phase exit rates largely corresponded to S-phase entry rates (Fig. 2d), we noted that during regeneration, S-phase exit rates were mostly smaller than entry rates (Fig. 6d). Taken together, dual pulse labeling revealed a complex pattern of alterations in cell cycle progression with a notable uncoupling of S-phase entry and exit. Lower exit rates are consistent with an overall increase in cells residing in S phase during regeneration. These observations underscore that S-phase frequencies, commonly used as a marker for proliferation at steady-state, only partially reflect proliferation speed.

**Figure 6.**
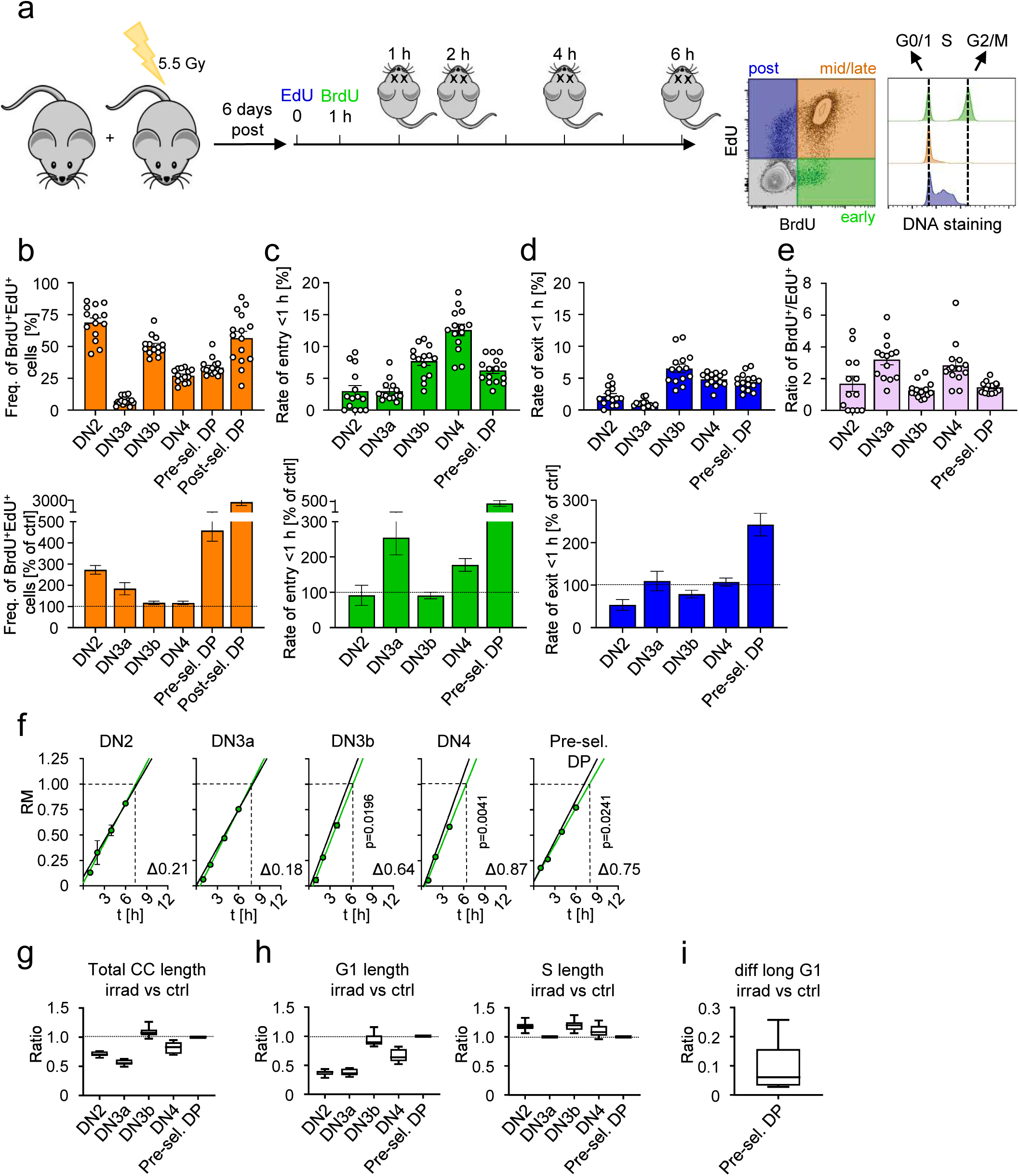
Population-specific cell cycle phase alterations determine thymus regeneration. **(a)** Schematic model of experiments. Ctrl and sublethally irradiated WT mice were administered EdU and BrdU 6 days post irradiation. Subsequent analysis including DNA content analysis was performed 1, 2, 4 or 6 h after the second pulse as described before. **(b-d)** Top: Frequencies of EdU^+^BrdU^+^ cells **(b)**, EdU^-^BrdU^+^ cells **(c)** and EdU^+^BrdU^-^ cells **(d)** of thymocyte subpopulations of irradiated mice, n = 14-15. Bottom: data is presented as % of ctrl (bottom row) with n = 16 ctrl mice and n = 14-15 irradiated mice. **(e)** Statistical analysis of irradiated WT thymocyte subpopulations to assess S-phase time duration based on RM values of BrdU^+^ cells (early S phase) over time (green dots). The green line represents a linear regression. The black line shows the corresponding ctrl. Numbers adjacent to linear regression show the difference in S phase time in h between ctrl and irradiated WT mice, with n = 4 ctrl mice and n = 1-4 irradiated WT mice for each time point. Analysis of significance between ctrl and irradiated WT mice was performed using unpaired t-test for the latest time point. **(f)** Box plots show total cell cycle length ratio of irradiated vs ctrl WT thymocyte subsets. **(g)** Box plots show G1- (left) or S-phase length (right) ratio of irradiated vs ctrl thymocyte subsets.

Lower exit than entry rates may be indicative of extended S-phase duration. To test this possibility, we analyzed dynamic cell cycle behavior *in vivo* using the previously established dual-pulse labeling approach in conjunction with DNA labeling to reveal S-phase progression of virtually synchronized cells over time. Estimated S-phase lengths in DN2 and DN3a thymocytes were similar with a trend towards longer duration during regeneration when compared to the steady-state (Fig. 6e). In DN3b, DN4 and pre-selection thymocytes S-phase duration was approximately 45 min longer upon regeneration. Thus, direct analysis of S-phase lengths supports the notion that, despite overall reduced cycling times, S-phase duration is increased in a model of post-irradiation thymus regeneration. Frequencies of single labeled cells within the post-selection DP subset were too low for a robust quantitative analysis. Next, we applied the ABM described above to model alterations in cycling behavior at the individual stage level. Consistent with the experimental observations, mathematical modeling revealed shorter overall cell cycle durations for DN2, DN3a, DN4 and pre-selection DP cells upon regeneration, with DN4 cell cycle duration being close to the steady-state situation (Fig. 6f). cell cycle length of DN3b was unaltered in regenerating thymi when compared to the steady-state. Decreases in cell cycle length could be completely attributed to a reduction in G1-phase duration for all populations within the population of actively cycling cells with the exception of pre-selection DP thymocytes, whereas S-phase duration was modestly increased in DN2, DN3b and DN4 thymocytes and remained unaltered in DN3a and pre-selection DP cells (Fig. 6g). We observed a substantial reduction in the fraction of ‘long G1’ non-cycling cells in pre-selection DP thymocytes, thus reconciling experimental data of an increased fraction of cycling cells in pre-selection DP cells without need of an increase in cell cycle speed as suggested by the mathematical model (Fig. 6i). We conclude that thymic regeneration post sublethal irradiation at cell cycle phase resolution is predominantly driven by increased rates of entry into cycle and shortening of the G1 phase, but not the S phase, predominantly in thymocyte subsets characterized by slow turnover at steady-state.

Thus, our studies reveal fundamental differences in how thymocytes adopt developmental stage dependent modulation of cell cycle length during steady-state T-cell development and perturbation-induced adaptation of cell cycle speed within populations during the regenerative process.

## Discussion

Studies on T-cell developmental dynamics in the thymus over the past decades have provided information on residence times, proliferation rates, and death during phases of selection, within individual thymocyte populations^12,13,15^, for review see^17^. Together, these data help to draw conclusions about division rates and overall cell cycle durations, but not about the duration of individual cell cycle phases. Such knowledge would constitute critical information to better understand regulatory principles of proliferation dynamics at steady-state and during therapy-induced regeneration. Thus far, analyses at cell cycle phase resolution were largely restricted to *in vitro* models e.g., employing FUCCI-transgenic systems in conjunction with time-lapse microscopy. Here, we have overcome this knowledge gap by devising experimental tools that allowed us to quantitate T-cell developmental dynamics at the cell cycle phase level. Furthermore, we have created an ABM to integrate all experimental information in order to generate testable hypotheses on cell cycle heterogeneity.

We accurately determined cell cycle phase durations in thymocytes of many distinct developmental stages at steady-state. Transition between slowly and rapidly cycling thymocyte populations was characterized by simultaneous contraction or expansion of G1 and S phases, supporting a so-called ‘stretch model’ of cell cycle regulation. This type of cycling behavior was first described to explain acceleration of B- and T-cell proliferation upon activation^24^. Similar to the situation of lymphocyte activation, thymocytes rapidly expand following β-selection from 4 x 10^5^ to 10^8^ cells, corresponding to 7 divisions within a period of only 2 – 4 days^15,17^. Contraction of both G1 and S phase might constitute the optimal solution to accommodate faithful DNA replication along with a minimal G1 phase. Indeed, in some DN3b cells our data and the ABM suggested a *bona-fide* absence of a G1 phase, similar to cell cycle patterns observed in embryonic stem cells^25^. In line with a previous study, the G2/M phase duration was uncoupled from stretching and contraction^26^. It remained essentially constant and its short duration of less than 1.5 h indicated that thymocytes immediately transit from DNA replication to mitosis without going through a sizable G2 phase.

Despite substantial numbers of cells in a prolonged state of cell cycle pause, especially in populations rearranging TCR loci (DN3a and pre-selection DP), FUCCI G1 reporter protein levels suggested that these cells did not adopt a state of quiescence (G0) comparable to hematopoietic stem cells. In line with an earlier study, a FUCCI-based quiescent state first emerged in DP thymocytes following selection, with quiescent cells accumulating in single-positive populations prior to thymic egress^19,27^. Although quiescence remains incompletely defined molecularly, a reduction in the expression of licensing factors, such as Cdc6 and Mcm2 and an increase in p27^kip^^1^ in DP thymocytes, supports our conclusion^28^. Re-entry into the cell cycle requires longer intervals than passing through a prolonged G1 phase. We suggest that a prolonged G1 duration is more compatible with a developmentally programmed trajectory, whereas quiescence is likely to be more compatible with naïve cells lying in wait for cognate antigen for undefined periods of time.

We noted distinct patterns of cell cycle heterogeneity. Heterogeneity in DN3b cells was best explained by varying G1-phase durations. In contrast, DN3a and pre-selection DP cell cycles were best explained including a subpopulation with a prolonged G1 phase that is not actively cycling within the time frame of the experiment. It is plausible that these subpopulations comprise cells undergoing TCR gene rearrangements, whereas actively cycling populations may comprise cells transitioning into or out of the respective subsets. In fact, a recent scRNAseq study has demonstrated that upon entry into the DP stage, thymocytes complete exactly one round of division before cycling ceases^29^. cell cycle heterogeneity was also apparent during longitudinal follow-up of cells and upon *in vitro* differentiation, substantiating the predictions derived from the ABM that variation is largely situated in G1 phase, whereas the transition from S to G1 is kinetically fixed in accordance with virtually constant durations of G2 and M phases. As the experimental system described in this study permits tracking of little more than one full cell cycle, it remains an open question whether the observed heterogeneity is a consequence of functional heterogeneity within phenotypically identical populations, in particular when cells undergo transitions between populations. Alternatively, cells may have unequal access to growth factors, such as IL-7 or display intrinsic stochasticity^30,31^.

To our surprise, thymocytes did not further increase cell cycle speed according to the stretch model during thymus regeneration. Rather, we observed substantially shorter G1 phases, whereas S phases remained constant or were even slightly extended. These findings are consistent with the possibility that feedback mechanisms controlling regeneration depend on extrinsic signals, to which cells preferentially respond during G1 phase. In contrast, our data suggest that S-phase duration is developmentally pre-programmed at the population level, but not critically affected upon perturbation. Furthermore, the activation of DNA damage checkpoints following irradiation may also counteract further contraction of S phases^32^.

In the ABM, we have assumed that cells complete a cycle before exiting a population. However, to date it remains elusive whether thymocyte populations progress to the subsequent developmental stage at defined phases of the cell cycle and if so at which phase the transition occurs. Resolving this question is complicated by the fact that marker-defined population boundaries as defined in the paradigmatic DN-DP-SP thymocyte scheme do not at some point in DN2 (the DN2a > DN2b transition)^9–11^. Onset of somatic recombination can occur in DN2b at an interval from commitment that yet remains to be defined or in DN3a ^33^. Similarly, there is no obvious biological difference between DN3b and DN4 apart from proliferation age and this population might by 1 division even expand to early DP cells ^29^. Additional modeling settings could be compared, such as entry in a specific phase of the cell cycle, or exiting after a certain time in the population rather than completion of the cycle. With regard to the impact of population transitions on cell cycle dynamics, we have confirmed that the entry of cells at a random phase of the cell cycle or after mitosis did not affect subsequent cell cycle phase durations (Fig. S3_5).

Mathematical modeling has been previously used to extract complex quantitative properties of T-cell development from experimental data^16,34^. We have used an ABM approach comparable to a model capable of reproducing the proliferation speed of lymphocytes^22^. The ABM allowed us to control the heterogeneity of phase durations within each cell, at the expense of computationally expensive simulations. Other mathematical models created to study thymocyte proliferation have used Ordinary Differential Equations (ODEs), which simulate the number of cells in each phase over time^16^. ODEs (sometimes modeled as Markov Chains^35^) can sometimes give analytical formulas for pulse-labeling over time, but structurally impose an exponentially distributed duration of each phase, where some cells stay for implausible short times in a phase^36^ . In consequence, it is not directly possible to compare synchronous or heterogeneous phase durations with those models. Interpretation of ODE-based models has been challenging, as they sometimes predicted extremely short and sometimes extremely long cell cycles, which are difficult to align with experimental data^35,37^. These inconsistencies may be attributed to limited sets of experimental data as well as limitations in permitting intra-population heterogeneity. A recent study employed an ODE model focusing on total DP thymocytes only, which considered multiple progression steps through cycle phases, limiting the possibility for cells to complete a cycle in a few seconds, and incorporated a final step of quiescent cells^38^. Separation of DP thymocytes into pre-selection (undergoing somatic recombination) and post-selection subsets in our study makes it difficult to directly compare both approaches. Alternatively, age- or DNA-structured models that account for a distribution of age or DNA inside each cell phase, could be used^39–41^. While being deterministic, these types of models (that often reduce to delay differential equations), can include heterogeneity and, unlike ODE, can introduce minimum residence time in each cell cycle phase. This could be a valuable alternative approach to keep most levels of complexity of the agent-based model while being less computationally expensive and providing more analytical possibilities than ODEs. Like other modeling approaches, the ABM is only able to capture certain levels of biological complexity. We validated this model employing a two-population model taking into account inflow of labeled precursors.

Taken together, our studies provide a combined experimental and computational framework to accurately determine cell cycle phase durations *in vivo*, integrating intra-population heterogeneity. We propose that this combination is widely applicable across organ systems and disease conditions. Applying this setup to study intrathymic T-cell development at steady-state and during regeneration, our studies have revealed a broad and population-specific spectrum of cell cycle adaptation to proliferative requirements. These findings illustrate the need for refined cell cycle analysis in situations of loss of proliferation control, such as cancer, or regenerative processes, which warrant preferential recovery of select cell types.

### Limitations of this study

This study focuses on defining cell cycle phase durations at high resolution in murine thymocytes in vivo. Although it would be of great interest to obtain similar data for human T-cell development, technical limitations still preclude high-resolution cell cycle phase analysis in human thymocytes. In the future, organoid technologies that more faithfully recapitulate developmental dynamics may enable such studies. The ABM described here in combination with experimental data is optimized to define cell cycle dynamics at high resolution. It still remains a challenge to simulate T-cell development in all its complexity in an all-encompassing model. In the ABM, we have used a pool of bystander cells in long G1. The longer the time-window, more continuous properties may add discrepancies between model simulations and experimental data, such as a continuous distribution of extremely long G1 durations within those long G1 cells, or the entry of new labeled cells into the long G1 pool. During regeneration after sublethal irradiation, thymocyte populations still dynamically change in size. Thus, technically non-steady-state conditions apply and this might influence the dynamics of labeling. Generally, our model is applicable to in vivo labeling of steady-state populations. Nevertheless, at the time scale of one cell cycle, population sizes are not substantially different and were considered as bona-fide steady-state for the purpose of this analysis. Further adaptations to account for non-equilibrium effects in rapidly growing populations would be particularly interesting for the field of cancer biology.

## Methods

### Key Resources Table

| REAGENT or RESOURCE | SOURCE | IDENTIFIER |
| --- | --- | --- |
| **FACS antibodies (clone)** |  |  |
| Monoclonal anti-BrdU antibody (3D4), FITC | BioLegend | Cat#364104;<br>RRID: AB_2564481 |
| Monoclonal anti-mouse CD4 (GK1.5), PE-Cy7 | BioLegend | Cat#100422;<br>RRID: AB_312707 |
| Monoclonal anti-mouse CD4 (GK1.5), PerCP-Cy5.5 | BioLegend | Cat#100434;<br>RRID: AB_893324 |
| IgM anti-mouse CD4 (RL1.72) | Own production | <sup>42</sup> |
| Monoclonal anti-mouse CD8α (53-6.7), BV510 | BioLegend | Cat#100751;<br>RRID: AB_2561389 |
| Monoclonal anti-mouse CD8α (53-6.7), PE-Cy7 | BioLegend | Cat#100722;<br>RRID: AB_312761 |
| IgM anti-mouse CD8 (M31) | Own production | <sup>43</sup> |
| Monoclonal anti-mouse CD25 (PC61), BV421 | BioLegend | Cat#102034;<br>RRID: AB_11203373 |
| Monoclonal anti-mouse CD25 (PC61), PerCP-Cy5.5 | BioLegend | Cat#102030;<br>RRID: AB_893288 |
| Monoclonal anti-mouse CD28 (E18), PE | BioLegend | Cat#122010;<br>RRID: AB_604078 |
| Monoclonal anti-mouse CD44 (IM7), APC-Cy7 | BioLegend | Cat#103028;<br>RRID: AB_830785 |
| Monoclonal anti-mouse CD44 (IM7), PerCP-Cy5.5 | BioLegend | Cat#103032;<br>RRID: AB_2076204 |
| Monoclonal anti-mouse CD69 (H1.2F3), PE | BioLegend | Cat#104508;<br>RRID: AB_313111 |
| Monoclonal anti-mouse CD69 (H1.2F3), APC | BioLegend | Cat#104514;<br>RRID: AB_492843 |
| Monoclonal anti-mouse CD71 (RI7217), APC | BioLegend | Cat#113819;<br>RRID: AB_2728134 |
| Monoclonal anti-mouse CD117 (2B8), APC-Cy7 | BioLegend | Cat#105826;<br>RRID: AB_1626278 |
| Monoclonal anti-mouse TCRβ (H57-597), BV421 | BioLegend | Cat#109230;<br>RRID: AB_2562562 |
| Monoclonal anti-mouse TCRβ (H57-597), APC-Cy7 | BioLegend | Cat#109220;<br>RRID: AB_893624 |
| Monoclonal anti-mouse TCRβ (H57-597), PE-Cy7 | BioLegend | Cat# 109222;<br>RRID: AB_893625 |
| **Chemicals** |  |  |
| DAPI (4',6-Diamidino-2-Phenylindole, Dilactate) | BioLegend | Cat#422801 |
| Bromodeoxyuridine (BrdU) | BioLegend | Cat#423401 |
| DNase I from bovine pancreas | Sigma-Aldrich | Cat#D4513 |
| EdU | Thermo Fisher Scientific | Cat#A10044 |
| Low-Tox®-M Rabbit Complement | Cedarlane | Cat#CL3051 |
| Lympholyte-M | Cedarlane | Cat#CL5031 |
| Permeabilization Buffer Plus | BD Biosciences | Cat#561654 |
| **Critical Commercial Assays** |  |  |
| Click-iT™ Plus EdU Alexa Fluor™647 Flow Cytometry Assay Kit | Thermo Fisher Scientific | Cat#C10635 |
| Fixation/Permeabilization Solution Kit | BD Biosciences | Cat#554714 |
| **Experimental Models: Organisms/Strains** |  |  |
| Mouse: C57BL/6J (WT) | Janvier Labs/ Charles River | N/A |
| Mouse: FUCCI (B6;Cg-Tg(FucciSG2M)<sup>#474Bsi</sup>-Tg(FucciG1)<sup>#639Bsi</sup>) | RIKEN | RBRC02704, RBRC02709 |
| **Software and Algorithms** |  |  |
| GraphPad Prism 7 | GraphPad Software | https://www.graphpad.com/ RRID: SCR_002798 |
| FlowJo 10™ | BD Biosciences | https://www.flowjo.com/solutions/flowjo RRID: SCR_008520 |

### Resource availability

#### Lead contact

Further information and requests for resources and reagents should be directed to and will be fulfilled by the Lead Contacts, A. K. or P. A. R.

#### Materials availability

This study did not generate new unique reagents.

#### Data and code availability

The code for the mathematical model is available at https://gitlab.com/haumeapoho/thypouille provided together with the Moonfit package^44^. The raw data used for simulations is provided within the code repository above.

### Experimental model and subject details

#### *In vivo* animal studies

C57BL/6J (WT) mice were purchased from Janvier Labs or Charles River. FUCCI mice were bred at the ZFE, Goethe University Frankfurt and at the ZVTH, Justus-Liebig-University Giessen. For all experiments, male and female mice were used between 8-12 weeks of age. All animal experiments were performed in accordance with local and institutional guidelines.

### Method details

#### Flow cytometry

Monoclonal antibodies specific for CD4 (GK1.5), CD8α (53-6.7), CD25 (PC61.5), CD28 (E18), CD44 (IM7), CD69 (H1.2F3), CD71 (RI7217), CD117 (2B8) and TCRβ (H57-597) were used conjugated to Brilliant Violet (BV) 421, phycoerythrin (PE), peridinin chlorophyll protein-Cy5.5 (PerCP-Cy5.5), PE-Cy7, Allophycocyanin (APC) or APC-Cy7 and were purchased from BioLegend. Cells were acquired using a BD FACSCanto II (BD Biosciences) and data was processed using FlowJo software (BD Biosciences). For data analysis, doublets and cells in sub-G0/G1 phase were excluded. For all panels, cells were defined as ETPs (CD117^hi^CD25^-^CD44^+^), DN1 (CD25^-^CD44^+^), DN2 (CD25^+^CD44^+^), DN2a (CD117^hi^CD25^+^CD44^+^), DN2b (CD117^lo^CD25^+^CD44^+^), DN3a (CD25^+^CD44^-^CD28^-^ or CD25^+^CD44^-^CD71^-^), DN3b (CD25^+^CD44^-^CD28^+^ or CD25^+^CD44^-^CD71^+^), DN4 CD25^-^CD44^-^CD28^+^, pre-selection DP (CD4^+^CD8^+^CD69^-^TCRβ^-^), post-selection DP (CD4^+^CD8^+^CD69^+^TCRβ^+^), SP4 (TCRβ^+^, CD4^+^) and SP8 (TCRβ^+^, CD8^+^).

#### Cell preparations

Thymi were meshed through a 70 µm cell strainer (Corning) to obtain single-cell suspensions. Cell numbers were determined using a CASY Cell Counter and Analyzer Model TT (Innovatis).

#### Dual pulse labeling using EdU and BrdU

For analysis of nucleoside analogue incorporation, mice were first intravenously injected with 1 mg EdU (Thermo Fisher Scientific) followed by 2 mg BrdU (BioLegend) 60 min later. Then, thymi were harvested at indicated time points. Single-cell suspensions were stained with monoclonal antibodies followed by EdU and BrdU staining procedures according to the manufacturer’s instructions using the Click-iT™ Plus EdU Alexa Fluor™ 647 Flow Cytometry Assay Kit (Thermo Fisher Scientific), BD Cytofix/Cytoperm™ (BD Biosciences), Permeabilization Buffer Plus (BD Biosciences) and treatment with DNase I from bovine pancreas (Sigma-Aldrich). To detect BrdU, samples were subsequently stained with FITC-conjugated anti-BrdU antibody (3D4, BioLegend). DNA content was analyzed by staining cells with DAPI (BioLegend).

#### Thymus regeneration

WT mice were sublethally irradiated (5.5 Gy). 6 days post irradiation, EdU/BrdU dual pulse labeling was performed as described before.

### Quantification and statistical analysis

All statistical analysis was performed using GraphPad Prism 7 software. Statistical parameters including number of mice (n) and number of replicates are described in the figure legends. Data are represented as mean plus or minus SEM. To compare ratios of two groups, data is presented as % of control and error bars indicate SE of ratios calculated as 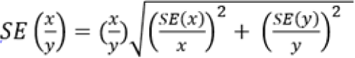. Analysis of significance between 2 groups of mice was performed using unpaired t-tests unless otherwise specified in the figure legends. For comparison between more than two groups, ordinary one-way analysis of variance (ANOVA) followed by Tukey’s test was used unless otherwise specified in the figure legends. P<0.05 was considered as significant.

### Agent-based model

#### General Strategy

*in vivo* EdU and BrdU labeling is a complex process since cells only receive labeling during the S phase, whose intensity depends on the speed of DNA incorporation (i.e., speed of the S-phase) and the duration of bioactive labeling. Further, mitosis causes a decrease of label intensity and doubles the amount of labeled cells. Therefore, a quantitative mathematical analysis is required. We designed a mathematical model to assess the likelihood *(cost)* of cycle phase durations *(parameters)* to reproduce the experimental kinetics of EdU/BrdU labeling *(observed variables)* as in the presented workflow (Fig. 4a). The model simulates a population of dividing cells *(agents)* and returns the expected labeling kinetics under a set of parameter values (in particular, the duration of cell cycle phases). The extent of deviation between simulation and the experimental data-points defines the objective ‘cost value’ to be minimized. Search heuristics that test large numbers of parameter values, allow finding cell cycle durations generating the minimal cost *(parameter optimization)*, and return the most likely phase durations to explain the experimental dataset. In the model, we simulate a population of dividing cells (the agents). We simulate separately the DN2, DN3a, DN3b, DN4 and pre-selection DPs with their own set of parameters (see Table S2). We explain how a simulation is performed from a hypothetical set of parameter values and how cells are updated over time for cell cycling, death and exit.

#### Cycle phases

The time evolution of the cell cycle is defined by a time-distribution for each possible event. The duration of the G1, S and G2/M phase are defined by three lognormal laws, with mean and standard deviations (in linear scale) named ***μG_1_, σG_1_, μS, σS, μG2/M, σG2/M***. Phases that are assumed non-variable are modeled with null standard deviation. Death is assumed to happen with a constant rate and therefore follows an exponential distribution of parameter **λ**. **λ** can be converted into a death rate **δ** per hour using **δ** = 1 / **λ**. A cell dies only if its time of death falls before it would perform mitosis.

#### Cells

Two types of cells are considered. Non-cycling cells contain either quiescent cells that stay in the G0 phase or bystander ‘long G1’ cells that do not cycle during the time of experiment, while cycling cells can be in G1, S and G2/M phases. Mitosis is included inside the G2 phase because it is fast and hardly distinguishable from G2. During a simulation, each cell carries information about the timing of its own past and future events (Table S3): time of birth (in G1 phase), ending time of each cycle phase, and time of death. A cell is also tagged with its generation and its instant DNA amount (labeled or unlabeled with EdU or BrdU).

#### Initialization of new cells

When a cell is created *(Algorithm 1)*, all its events are sampled according to the time-distributions for the cycle phases and death. This assumes independence between cells: that their cell cycle is predefined at birth and is not influenced by dynamical factors such as changes in population size. When daughter cells are created by mitosis, the current time becomes their time of birth, and each following event for both cells are sampled once for all, from the respective death or phase duration distributions. At the start of a simulation, in order to start with a steady-state population of cells in each stages of the cell cycle, cells are generated in the middle of the cell cycle: each event is sampled and shifted to the past, such that the current time falls randomly inside one of the phases proportionally to their duration. Finally, a quiescent cell is created by setting its state to G0 and giving ***‘+∞’*** as a time point for all phase progression events so they never happen.

#### Population initialization

A population of size ***Ncells*** is simulated, with a fraction of non-cycling cells noted ***q0***. We assume that EdU or BrdU injections do not significantly alter proliferation during the time frame of the experiment, and that non-cycling cells are constant over time. Previous mathematical models for EdU labeling assumed an exponential population growth without exit^35,37,38,45^, which is a valid assumption in cell cultures and simplifies the mathematical formulation but might introduce biases in the present case. As in some population models of thymic development^46–48^, we instead assume the cells stay in a population for an average number of divisions ***Ndiv*** before they exit, coupled with an inflow of unlabeled progenitors to maintain population time. For instance, a value of ***Ndiv = 1.6*** means that 40% of cells will perform one division whereas 60% perform two divisions, and ***Ndiv – floor(Ndiv)*** defines the fraction of cells performing one more division.

At equilibrium, each generation contains more cells than the previous generation (provided death is low and the population is expanding between inflow and outflow). Therefore, if a population of cells is generated by uniformly sampling generations among cells, this population is not stable over time. *Algorithm 2* shows how to generate an initial pool of cells with appropriate generations. The function ***equilibriumGenerations*** calculates the distribution between generations at equilibrium. The average time to complete a full cycle ***T*** is derived from each cycle phase time distribution, and the death rate **δ** from the death time distribution. During a period of time ***T***, the cells will expand on average 2 times by proliferation and die ***T.*δ** (approximation of ***1 – exp(-t.* δ)**, provided death is low). Therefore, the average expansion rate of the population is ***X = 2(1- T* δ)**. The fraction of cells in each generation is calculated knowing that each generation is ***X*** times bigger than the previous one, and only a fraction of cells enter the last generation. The function **GenerateCells** then generates a population of ***Ncells*** cells, either non-cycling (fraction ***q0***) or at a random phase of the cycling from *Algorithm 1*, with appropriate generations.

#### Simulation of a dual pulse labeling

*Algorithm 3* explains how the population of cells are updated at each time-step at time ***time,*** updated every ***dt***, and assuming an instant level of bioavailability of EdU and BrdU. Since each cell contains the information of its future event, each cell ***t*** is checked for its events happening between ***time - dt/2*** and ***time + dt/2.*** Death is checked first and leads to cell removal. Alive cells are updated for their DNA level depending on their phase (DNA levels increase linearly from 1 to 2 during the S-phase, Fig. 4b). The newly synthesized DNA during this ***dt*** time is labeled proportionally to the current levels of EdU and BrdU (∈ [0,1] each). Phase completion events are performed by changing the cycle state. At the end of the G2/M phase, if ***t*** was not completing its last division, two new daughters are created at G1 with their own new events and an incremented generation compared to the mother cell, ***t***, which is then removed from the simulation. If the two cells are reaching the last possible generation, only with chance ***Ndiv – floor(Ndiv)*** they will stay and be tagged for exit after the next mitosis, or exit directly (be removed). Finally, a constant inflow of unlabeled cells is calculated as needed to maintain the ‘generation 0’ constant, ensuring the full population to be constant over time.

#### Identification of cell phase durations

All parameters necessary to define and perform a simulation are listed in Table S2. Phase durations and width are unknown and ‘fitted’ by parameter estimation within the given minimum and maximum boundaries, depending on the chosen hypothesis of phase heterogeneity (Fig. 4b). The list of fitted variables is shown in Table S4. The fraction of non-cycling cells ***q0*** is either fitted (simulations with long G1 cells) or taken as a background value of 4% taken as an average value from experimental data. Finally, we chose to simulate 10,000 cells per population for the sake of computational complexity. EdU^+^ or BrdU^+^ cells were defined as cells with more than 0.01 of labeled DNA (on a scale from 0 to 2). We assumed each pulse to be efficient for a duration of 45 minutes^49,50^.

Each time a simulation is performed (Algorithm 4), the cost function between simulation and data is the Mean Standard Error (MSE) normalized for each curve separately by its average value along all time points, such that each curve has a balanced contribution to the total cost. Due to the low number of mice per time point, we did not include experimental standard deviations inside the cost calculation, as it was putting the weight only on points with low standard deviation by chance, and the fittings were not of good quality.

We have used an evolutionary strategy algorithm as a search heuristic for parameter estimation, using the Moonfit framework in C++^44^. A population of possible parameter sets (individuals) is generated with uniform values within the given unknown parameter boundaries. Each individual also carries a mutation speed on its own for each parameter, and is associated with a fitness, that is, the cost of a labeling simulation performed with its parameters. New individuals are generated by mutation (changing parameter values) or recombination between two parents. Mutation followed a normal distribution sampled for all the parameters independently at the same time. The SBX crossover was used for recombination^44^, sampling of parents was made proportional to their fitness, defined as the competitive advantage relative to the current worst individual fitness. Offspring did not replace the parents, but only the best individuals (including parents and progeny) were kept after each round of mutations and recombination, to maintain the number of individuals. Each optimization was performed ten times separately, using a population of 250 individuals along 100 rounds of generation/selection. Every round, 20 % of the population was generated as new individuals by cross-over, while 50 % of the population size was generated by mutation. Therefore, one optimization tests 25,000 possible parameter sets, each simulating the labeling of a population of 10,000 cells.

#### Two-populations model

To simulate the effect of inflow of cells from the ancestor population of the main population of interest, the two-population model simulates the ancestor population with ***5 x Ncells*** cells and following previously identified cell cycle durations (i.e., none of the ancestor population parameters are part of the fitting). For each time-step dt, the main population requests a certain number of new inflow cells and picks randomly from the pool of cells that exited the ancestors at this same time-step (which was the reason to simulate a much bigger ancestor population to always have cells available). If not enough cells are available, an inflow deficit accumulates, and more cells are taken in the next time-steps. We have checked that such a deficit never accumulated in the simulations, and 5 x Ncells was sufficient as the ancestor population size. Contrarily to the one-population model, where cells were generated at any cycle phase, the ancestor cells only exit after mitosis and therefore always enter at the beginning of their G1 phase, which allows to have a consistency between entry at mitosis and exit at mitosis in the main population. However, we checked that entry at a random cell cycle phase from the ancestors to the main population, gave the exact same identified phase durations (Fig. S3_5), showing that the phase of entry in a population did not hamper our capacity to identify phase durations.

#### Prediction of cell cycle modulation after irradiation

To estimate if irradiation had an impact on different phases of the cell cycle, we simulated side-by-side the dual-pulse labeling of control (CTRL) and irradiated (IRR) mice, and optimized the summed cost of both simulations to the respective datasets. Of note, the CTRL dataset was generated together with the IRR dataset and comes from distinct experiments than the kinetic labeling used in the previous figures (WT dataset, see supplementary data). Different hypotheses for which cycle phase is impacted by irradiation are considered (Fig. S5_1). The null hypothesis (‘none’) assumes that CTRL and IRR mice have the same cell cycle phase durations and both simulations share all parameters. The ‘diff G1’, ‘diff S’, ‘diff G2/M’ assume only one phase is modulated by irradiation, and both CTRL and IRR simulations share all parameters except the duration of the respective phase. Finally, the ‘diff All’ condition assumes that all phase durations are impacted by irradiation and the two simulations have independent parameters. The condition in green boxes (Fig. S5_1) shows the minimum model complexity that explains the CTRL and IRR datasets with the best cost. In DN3a and pre-selection DPs, a different G1 duration between CTRL and IRR was necessary to explain the data, but not any other phase, suggesting that irradiation only modulated the G1 phase. In other populations, a modulation of each phase was necessary to explain the data.

#### Comparison of model hypotheses using AICc

For each model hypothesis (which cell cycle phase is variable between cells and whether long G1 cells are allowed), we calculated the corrected Akaike Information Criterion (AICc) which compares the cost of the best curve for each hypothesis, giving a penalty to hypotheses with more parameters. The lowest AICc describes the best hypothesis. The AICc is calculated with the following formula, where experimental data is assumed to follow a gaussian distribution, ***n*** represents the amount of independent fitted points (***n*** *= 44* for 4 time-points and ***n*** = 62 for 6 time-points), and k is the number of unknown parameters.

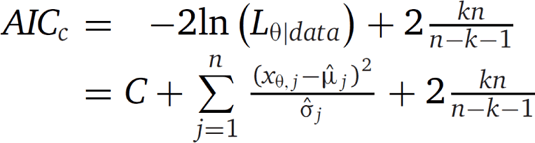

## Acknowledgments

We are grateful to Dr. Jonas Blume and Eleni Dapergola for technical help during the early phase of the project We are grateful to Julia Häusler and Bedriska Reitz for technical support. We thank Prof. Dr. Anja Hauser (DRFZ Berlin) for providing the FUCCI mouse line and Dr. Ines Kühnel for critical reading of the manuscript and helpful discussions. This work was funded by grants from the German Research Foundation (DFG), KR2320/6-1 and EXC62 ‘REBIRTH’ to A.K.

## Author contributions

1. H. K.-S., P. A. R., M. M.-H. and A. K. designed all experiments and analyzed data. H. K.-S. and Z. G. conducted all experiments. P. A. R. generated the mathematical model and performed simulations. N. A. V. analyzed data. H. K.-S., P. A. R., V. G., M. M.-H. and A. K. wrote the manuscript.

