## Supplementary Information for "High-resolution mapping of cell cycle dynamics during T-cell development and regeneration *in vivo*"

### Figure S1

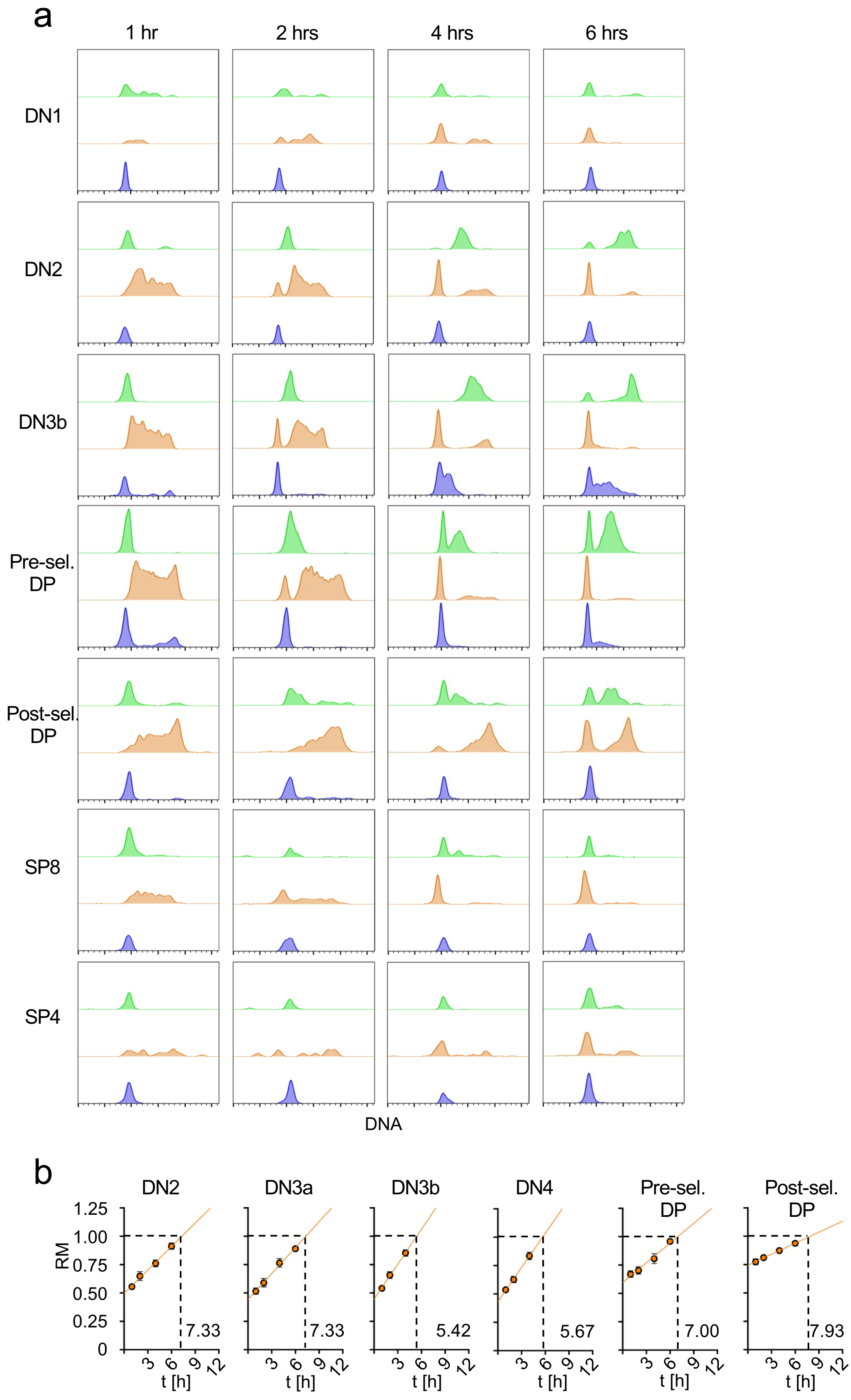


**Figure S1. (a)** Representative flow cytometric histograms of DNA content of DN1, DN2, DN3b, pre- and post-selection DP, SP8 and SP4 thymocytes of WT mice at indicated time points. Each plot depicts an overlay of the DNA content of $EdU^-BrdU^+$ (green), $EdU^+BrdU^+$ (orange) and $EdU^+BrdU^-$ (blue) cells. **(b)** Statistical analysis of WT thymocyte subpopulations to assess S-phase duration based on RM values of $EdU^+BrdU^+$ cells (mid/late S phase) over time (orange dots). The orange line represents the resulting linear regression. Numbers adjacent to linear regression show S-phase duration in h calculated based on linear regression, n = 3-5 mice for each point in time, data from 2 independent experiments.

### Figure S2

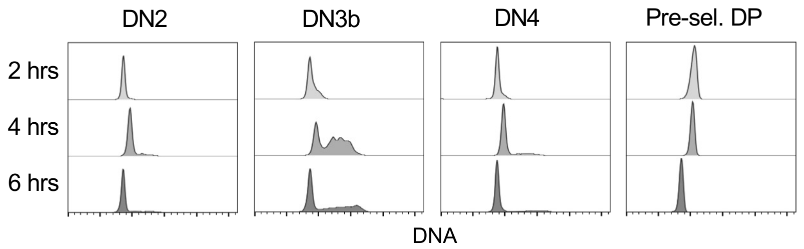


**Figure S2.** Representative flow cytometric histograms of DNA content of EdU$^-$BrdU$^-$ DN2, DN3b, DN4 and pre-selection DP thymocytes of WT mice over time. Each plot represents an overlay of the DNA content at 2, 4 and 6 h.

### Figure S3_1

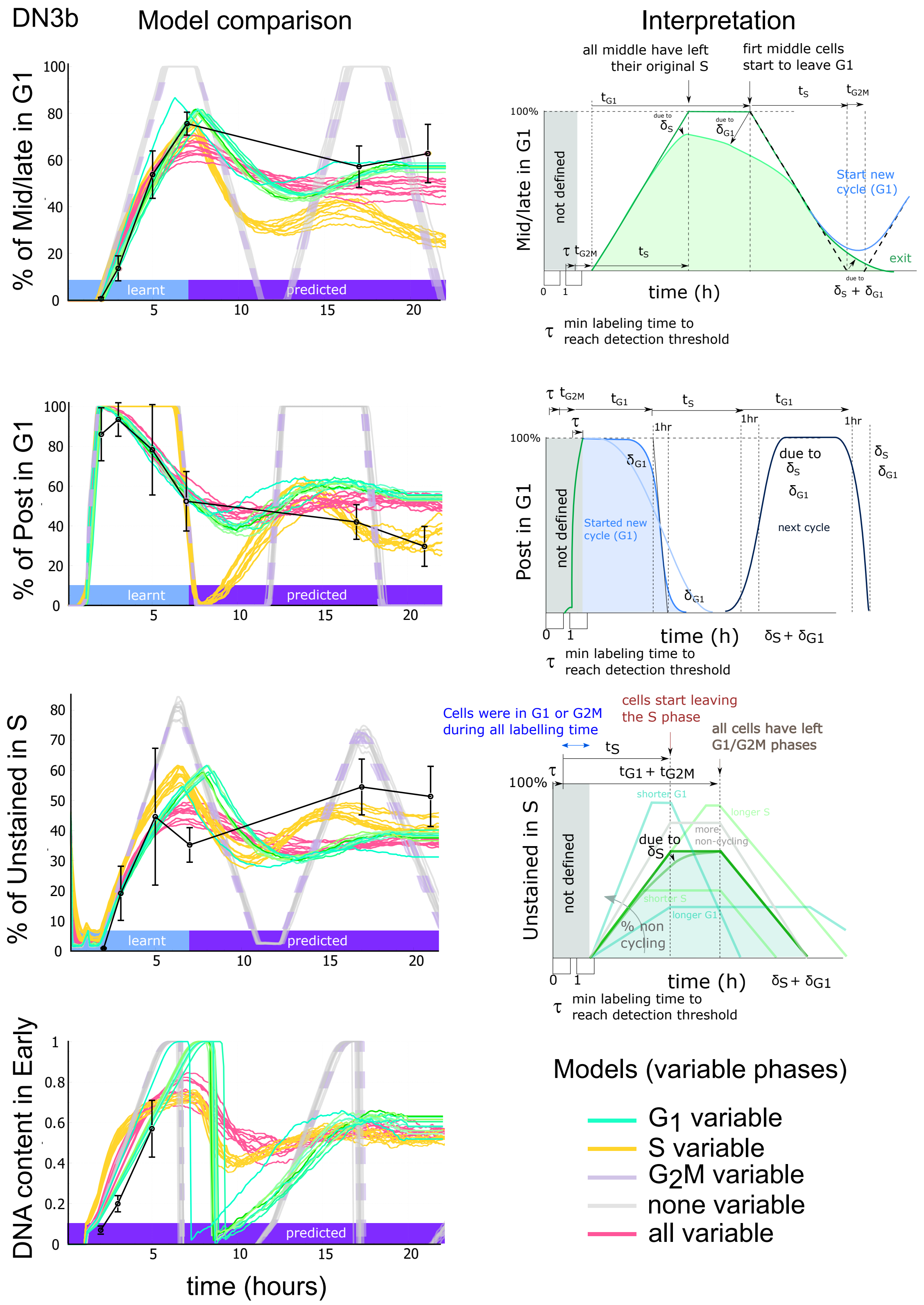


**Figure S3_1. Variation in G1-phase but not S-phase duration is required to explain the dual-pulse labeling kinetics of DN3b thymocytes.** Left: The simulation curves under the phase heterogeneity hypotheses are shown for the most informative experimental variables (see Figure S3_2 for comparison with all measured variables). Right: The interpretation of the curves is shown based on the average duration of each phase (tG1, tS and tG2M); the expected effect of each phase variation ($\delta$S, $\delta$G1 and $\delta$G2/M) and of the amount of "long G1 cells" (% of non cycling).

### Figure S3_2

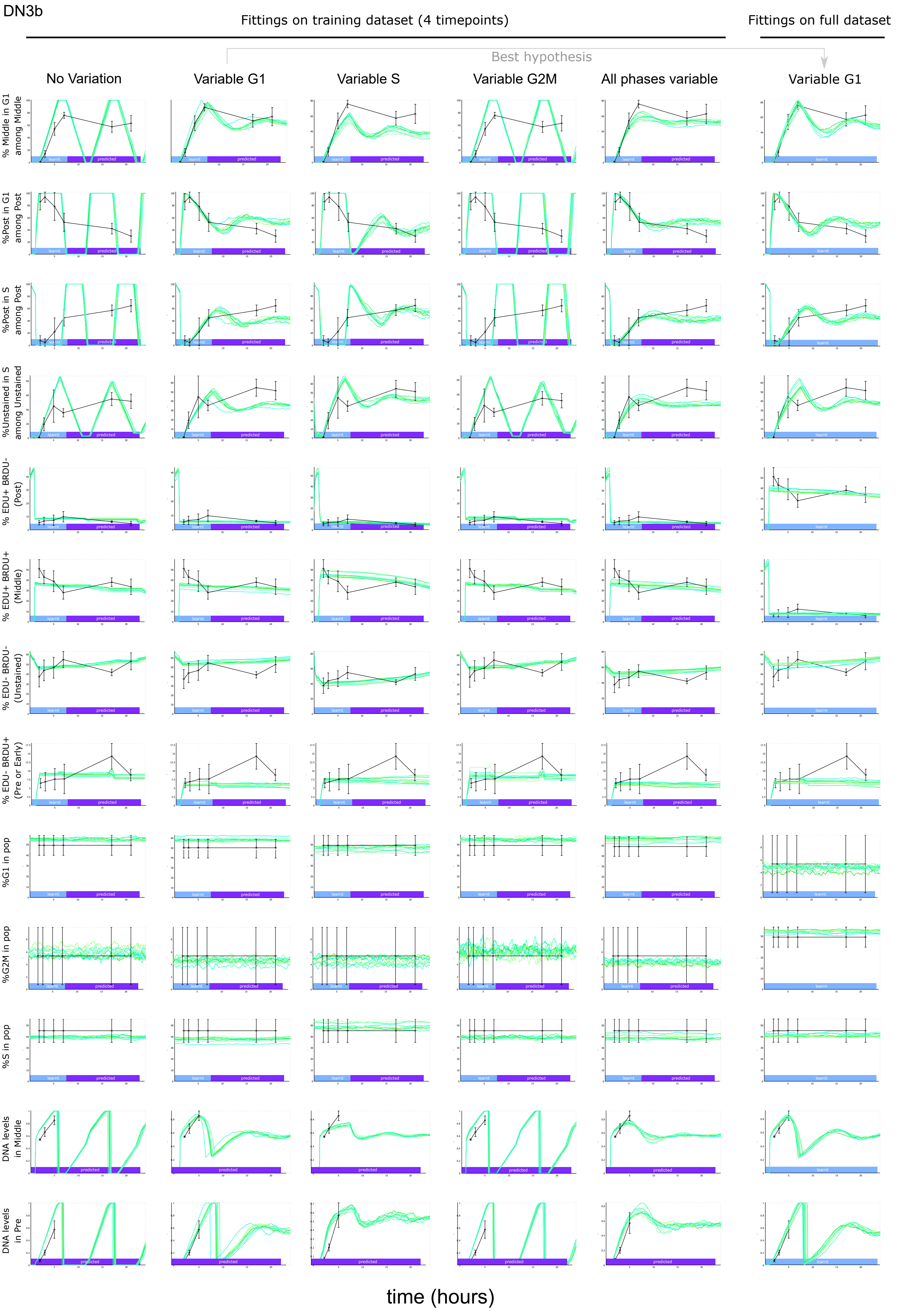

DN3b
Fittings on training dataset (4 timepoints)
Fittings on full dataset
Best hypothesis
No Variation
Variable G1
Variable S
Variable G2M
All phases variable
Variable G1
% Middle in G1 among Middle
%Post in G1 among Post
%Post in S among Post
%Unstained in S among Unstained
% EDU+ BRDU- (Post)
% EDU+ BRDU+ (Middle)
% EDU- BRDU- (Unstained)
% EDU- BRDU+ (Pre or Early)
%G1 in pop
%G2M in pop
%S in pop
DNA levels in Middle
DNA levels in Pre
learnt
predicted
time (hours)

### Figure S3_2

**Figure S3_2. Variation in G1-phase but not S-phase duration is required to explain the dual-pulse labeling kinetics of DN3b thymocytes (detailed curves).** The experimental data is shown in black, and the simulations from 10 independent fits are shown in green according to different model hypotheses on the stochasticity of phase durations at the population level. No variation: all the cells have the exact same duration of each cell cycle phase. Variable phases: only one phase is different between cells, and picked from a lognormal distribution. All phases variables: each phase follows a different log-normal distribution between different cells. For the top 11 observed variables, the 4 early time points are used for fitting (learnt), without knowledge of the remaining two time points (predicted). The bottom two variables, that depict the amount of DNA in $EdU^{-}BrdU^{+}$ and $EdU^{+}BrdU^{+}$ cells, were excluded from fitting and used as an independent qualitative validation dataset. No variation in phase durations, or only variation in the S phase cannot explain the dynamics of the $EdU^{+}BrdU^{-}$ cells or the DNA levels in $EdU^{-}BrdU^{+}$ cells, while variation in the G1 phase is sufficient to recapitulate all the observed variables. Allowing all phases to be variable does not improve the quality of the curves.

### Figure S3_3

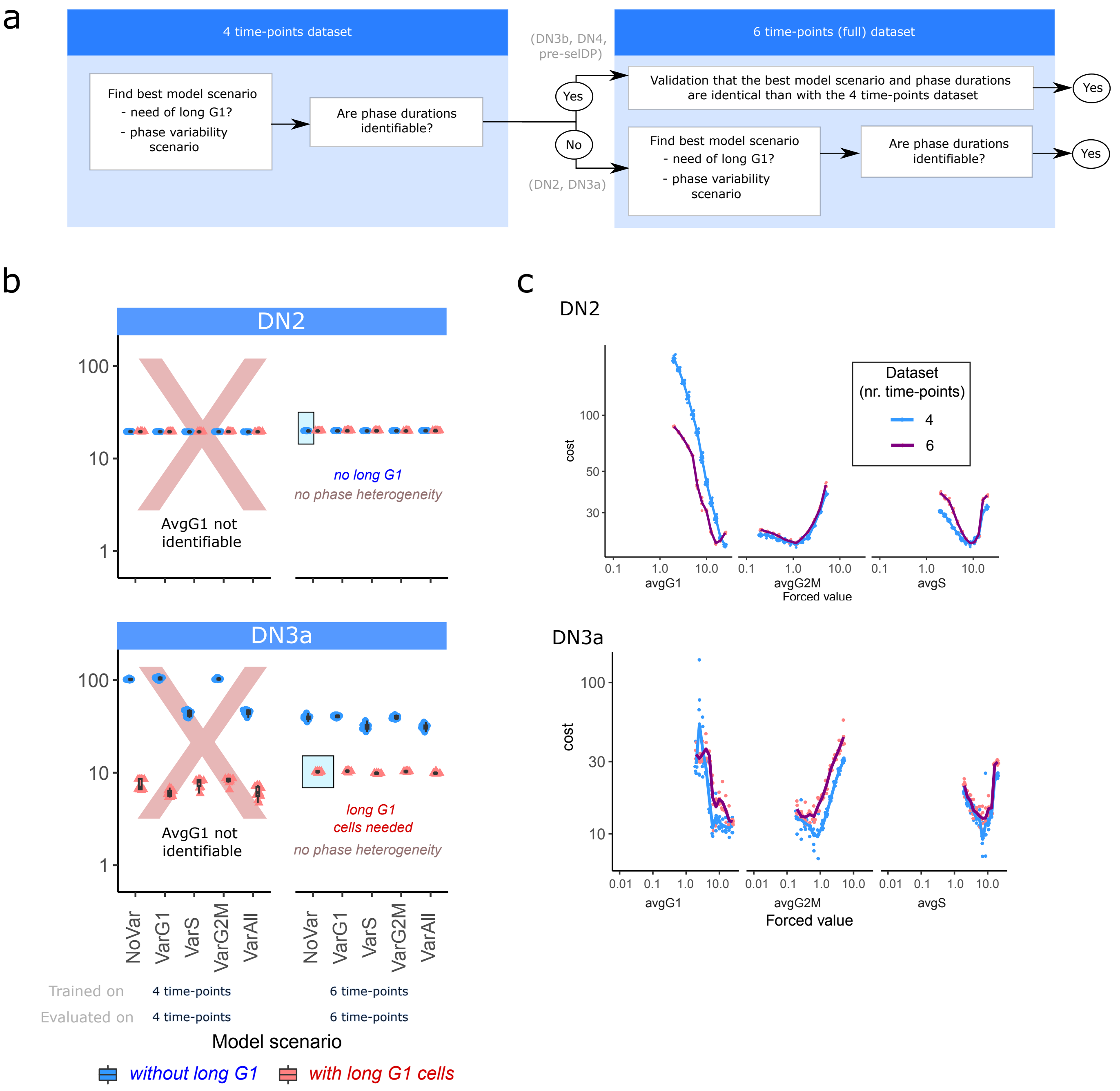


**Figure S3_3. Minimal amount of data necessary to identify cell cycle phase durations. (a)** Strategy to decide whether 4 or 6 time-points are necessary. **(b)** cost of cell cycle heterogeneity scenario in the presence (red) or absence (blue) of long G1 cells. For DN2, phase heterogeneity or long G1 cells do not improve the cost of simulations, meaning that DN2 labeling can be explained without phase heterogeneity or long G1 cells. **(c)** Identifiability analysis of phase durations with 4 (training dataset) or 6 (full dataset) time points. It is not possible to identify the G1 duration of DN2 and DN3a with only four time points.

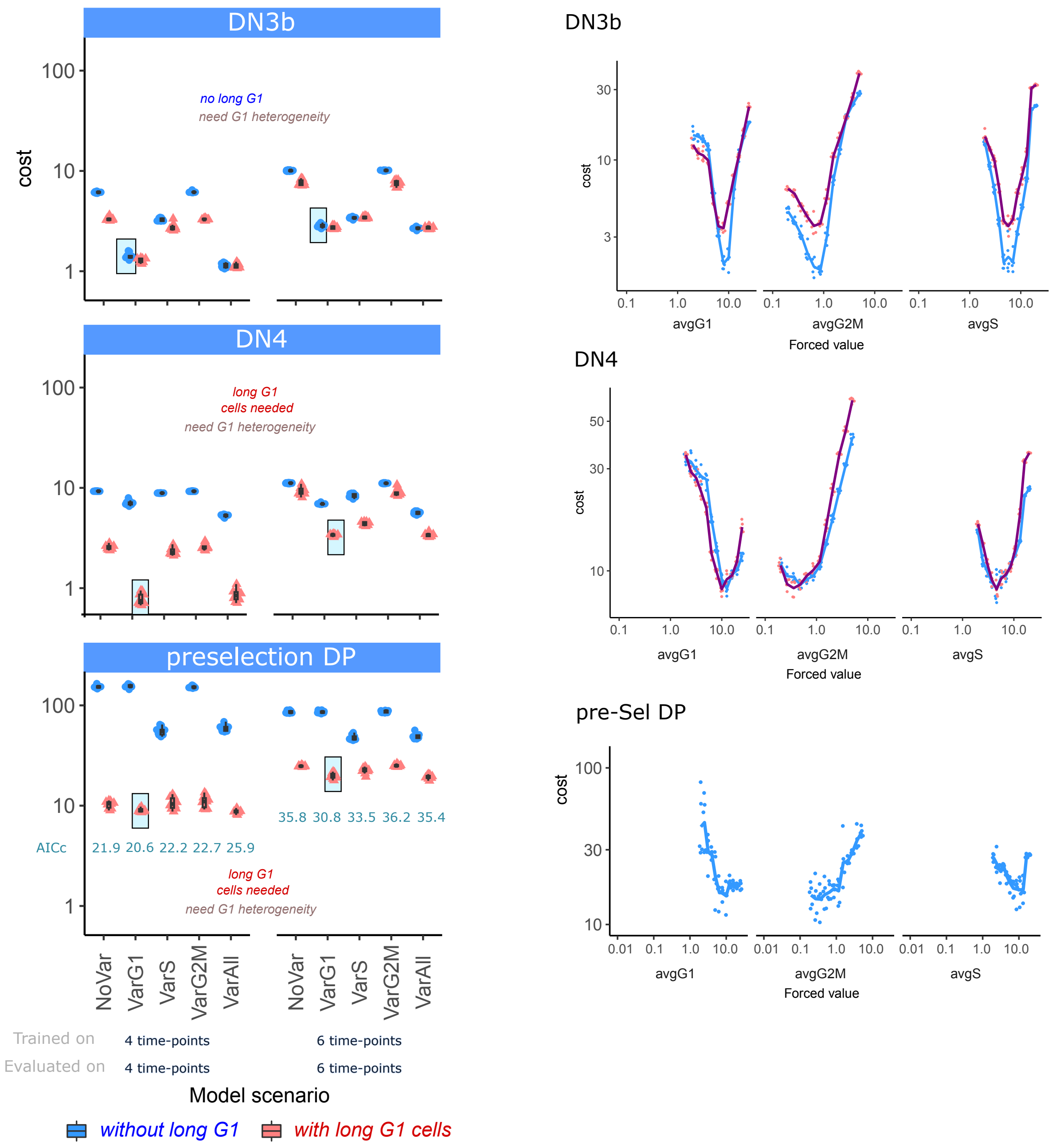


**Figure S3_3. (continued) (left)** cost of cell cycle heterogeneity scenario in the presence (red) or absence (blue) of long G1 cells. **(right)** Identifiability analysis of phase durations with 4 (training dataset) or 6 (full dataset) time points.

### Figure S3_4

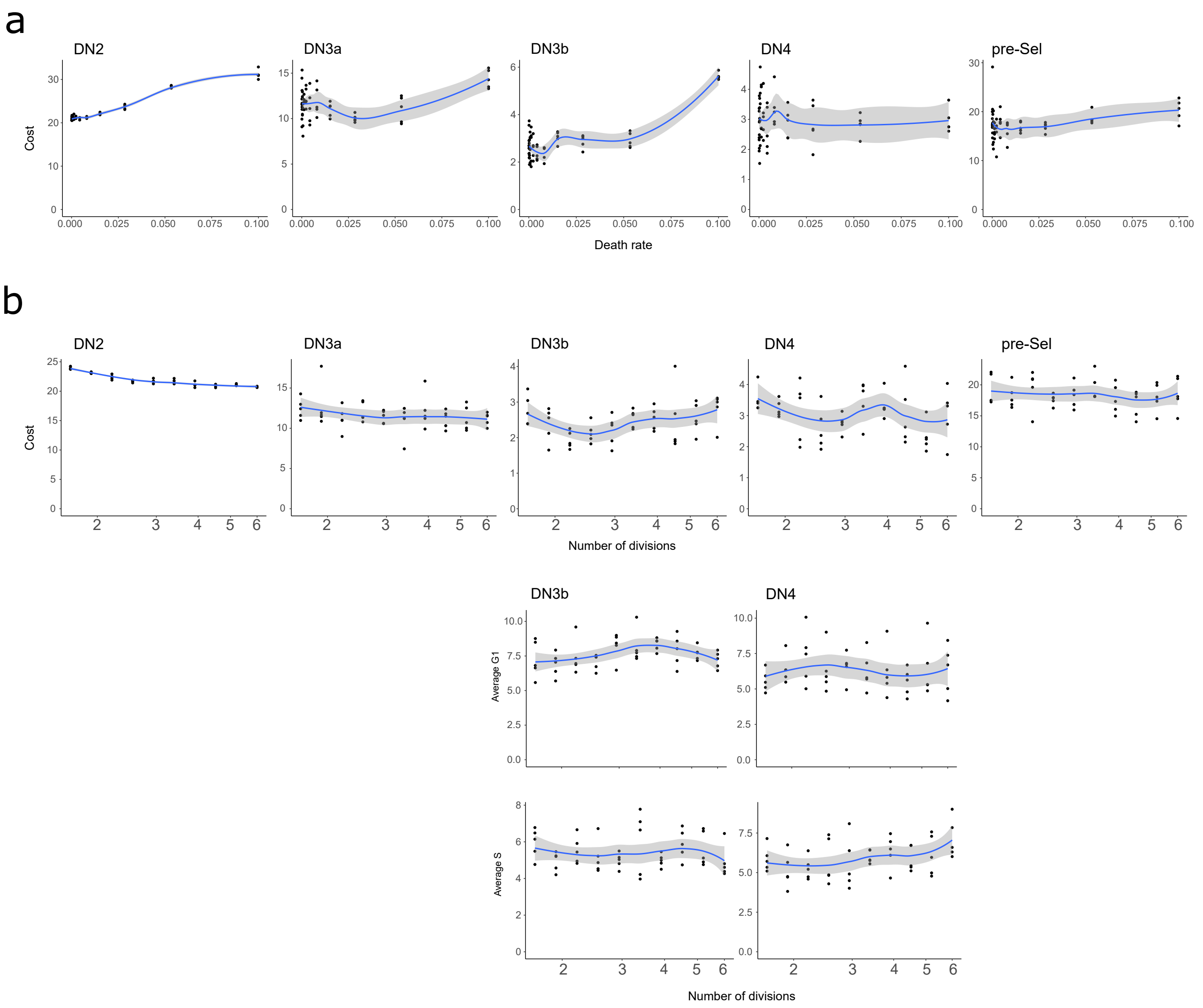


**Figure S3_4. Robustness of our phase duration identification to fixed parameters (number of divisions and death rate). (a,b)** Identifiability of the death-rate (a) and the number of divisions (b) by profile likelihood. **(c)** identified duration of G1 and S phases when for each fixed number of divisions.

### Figure S3_5

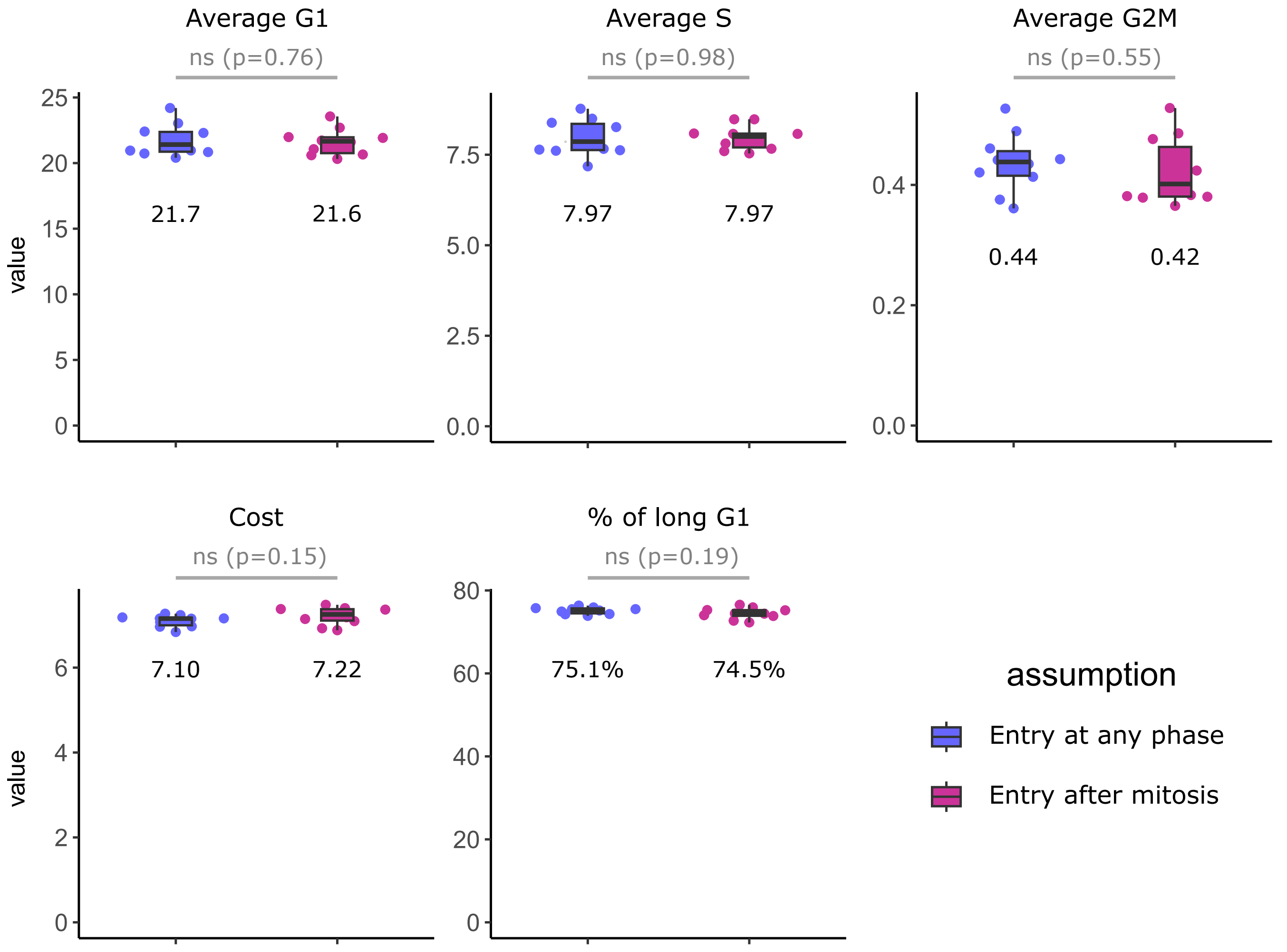


**Figure S3_5: The cycle phase at which cells enter a population do not impact the identification of cell cycle phase durations.** Comparison of phase durations using the two-populations model on the full dataset on the DN3a population, when cells transit from the ancestor to the main population at a random phase of the cycle, or just after mitosis.

### Figure S3_6

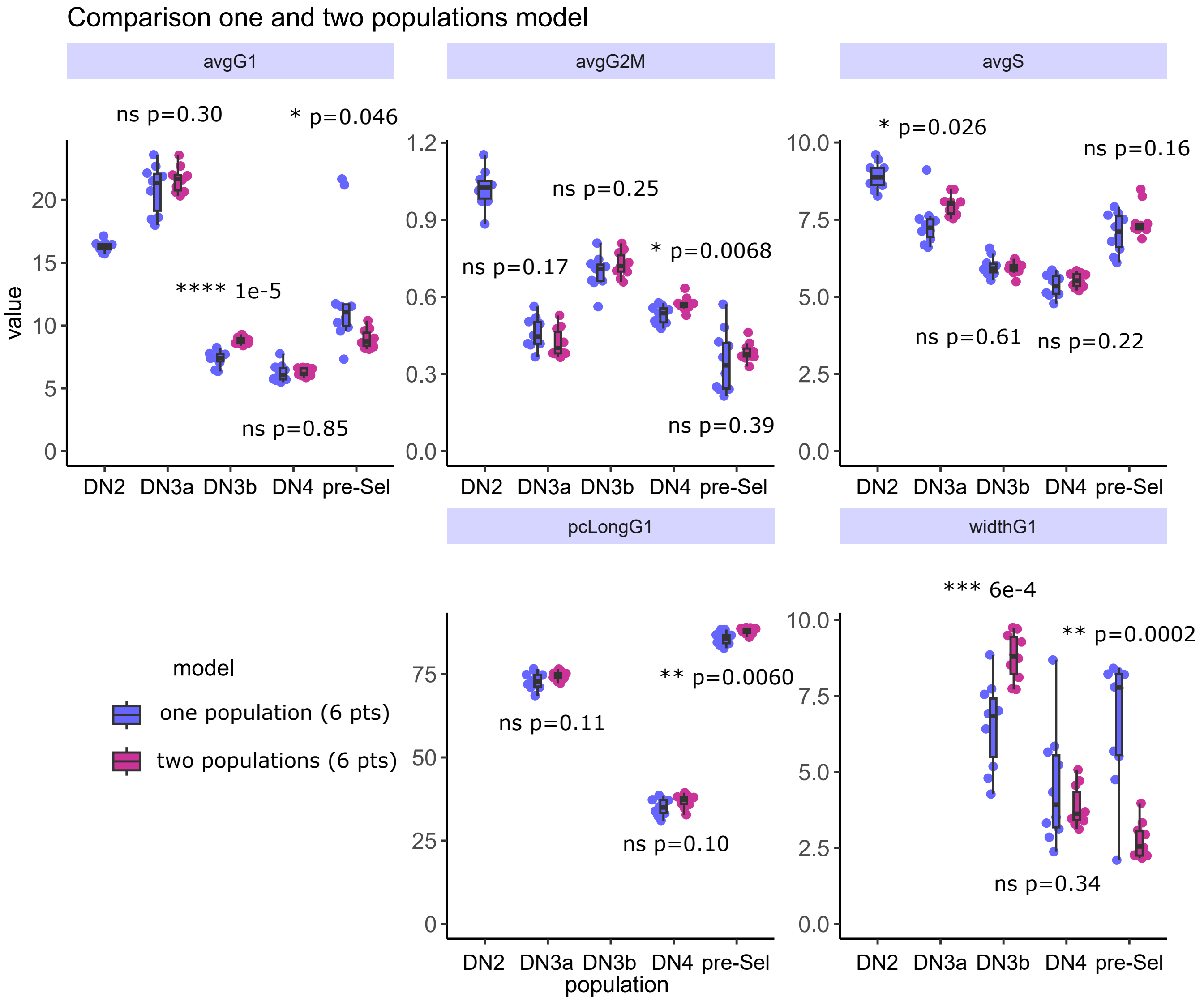


**Figure S3_6: The phase durations identified by the one-population and two-population model show minor differences,** showing that the biological complexity missing in the one-population model has only a minor impact on the identification of phase durations.

### Figure S4

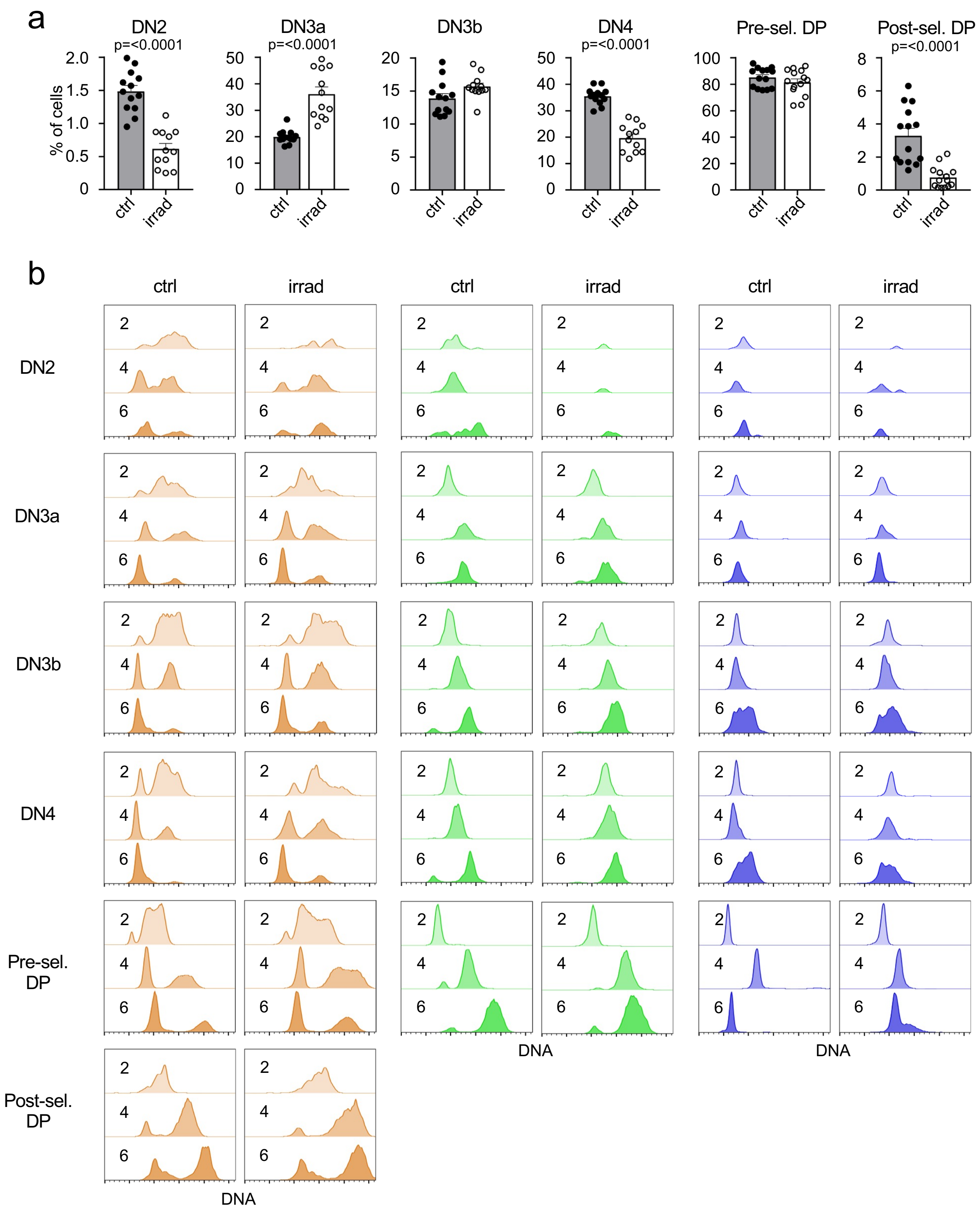


**Figure S4. (a)** Thymocyte subset composition from ctrl (grey, black dots) and irradiated WT (white, white dots) mice indicated as frequencies with n = 13-14 ctrl mice and n = 12-14 irradiated WT mice. **(b)** Representative flow cytometric histograms of DNA content of different thymocyte subsets of ctrl and irradiated WT mice over time of $EdU^+BrdU^+$ (orange), $EdU^-BrdU^+$ (green) or $EdU^+BrdU^-$ (blue) cells. Each individual plot represents an overlay of the DNA content at 2, 4 and 6 h.

### Figure S5

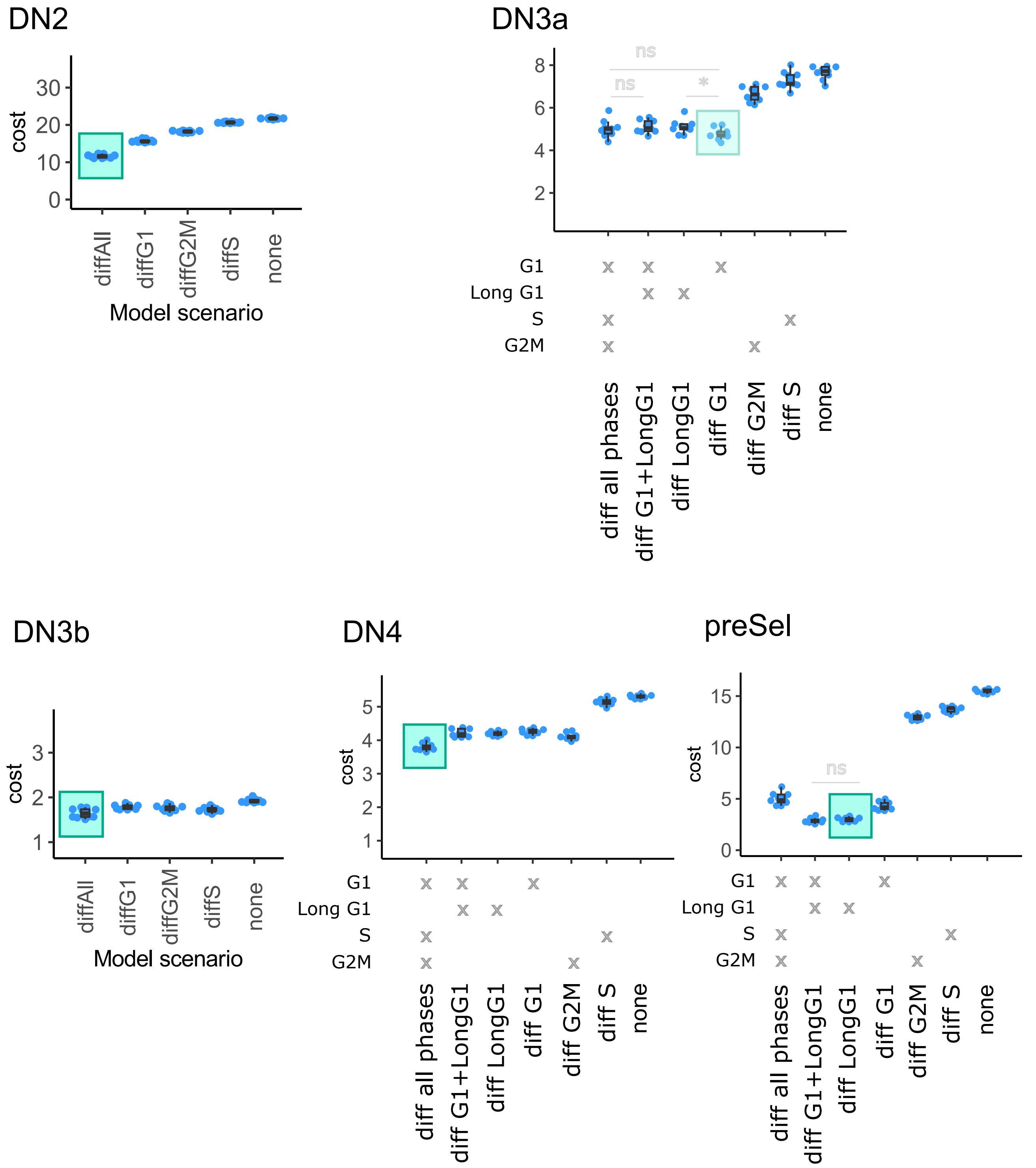


**Figure S5. Comparison of hypotheses of modulated phase duration due to irradiation.** For each population, cost of simulating the control (CTR) and irradiated )IRR) datasets together with the same phase durations except the phases hypothetically modulated by irradiation (x axis). The model with minimal complexity raising a minimal cost is kept and shown in a green box.

### Table S1

**Minimal — One-population**

| | Best Model | Long G1? | nec. dataset | cycle duration | G1 duration | S duration | G2M duration | % long G1 | width G1 cycle |
|---|---|---|---|---|---|---|---|---|---|
| DN2 | No var | N | full (6 points) | **27.4** [ 26.8 , 28.1 ] | **17.0** [ 16.6 17.5 ] | **9.4** [ 9.0 9.8 ] | **1.05** [ 1.0 1.1 ] | | |
| DN3A | No var | Y | full (6 points) | **28.7** [ 26.7 , 30.7 ] | **20.9** [ 19.0 , 22.8 ] | **7.4** [ 6.6 , 8.1 ] | **0.46** [ 0.4 , 0.5 ] | **73.0** [ 70.5 75.4 ] | |
| DN3B | Var G1 | N | train (4 points) | **15.8** [ 15.1 , 16.6 ] | **8.9** [ 8.5 9.3 ] | **6.2** [ 5.7 6.7 ] | **0.75** [ 0.7 0.8 ] | | **8.9** [ 8.2 9.6 ] |
| DN4 | Var G1 | Y | train (4 points) | **12.9** [ 11.5 , 14.2 ] | **6.6** [ 5.4 7.8 ] | **5.7** [ 5.4 6.0 ] | **0.61** [ 0.6 0.7 ] | **40.3** [ 36.7 43.8 ] | **5.3** [ 2.9 7.8 ] |
| preSel | Var G1 | Y | train (4 points) | **17.9** [ 15.8 , 20.1 ] | **8.1** [ 6.3 9.9 ] | **9.3** [ 8.0 10.7 ] | **0.5** [ 0.4 0.6 ] | **91.5** [ 90.2 92.8 ] | **3.7** [ 0.9 6.6 ] |

**Full data — One-population**

| | Best Model | Long G1? | nec. dataset | cycle duration | G1 duration | S duration | G2M duration | % long G1 | width G1 cycle |
|---|---|---|---|---|---|---|---|---|---|
| DN2 | No var | N | full (6 points) | **27.4** [ 26.8 28.1 ] | **17.0** [ 16.6 17.5 ] | **9.4** [ 9.0 9.8 ] | **1.05** [ 1.0 1.1 ] | | |
| DN3A | No var | Y | full (6 points) | **28.7** [ 26.7 30.7 ] | **20.9** [ 19.0 , 22.8 ] | **7.4** [ 6.6 , 8.1 ] | **0.46** [ 0.4 , 0.5 ] | **73.0** [ 70.5 75.4 ] | |
| DN3B | Var G1 | N | full (6 points) | **14.0** [ 13.2 14.9 ] | **7.4** [ 6.8 , 8.0 ] | **6.0** [ 5.7 , 6.3 ] | **0.70** [ 0.6 , 0.8 ] | | **6.6** [ 5.1 8.0 ] |
| DN4 | Var G1 | Y | full (6 points) | **12.1** [ 11.2 13.0 ] | **6.2** [ 5.6 , 6.9 ] | **5.4** [ 5.0 , 5.7 ] | **0.53** [ 0.5 , 0.6 ] | **35.1** [ 32.6 37.6 ] | **4.5** [ 2.6 6.4 ] |
| preSel | Var G1 | Y | full (6 points) | **20.0** [ 14.6 25.4 ] | **12.5** [ 7.7 , 17.4 ] | **7.1** [ 6.5 , 7.7 ] | **0.4** [ 0.2 , 0.5 ] | **85.7** [ 83.8 87.6 ] | **6.7** [ 4.6 8.8 ] |

**Full data — Two-populations**

| | Best Model | Long G1? | used dataset | cycle duration | G1 duration | S duration | G2M duration | fraction long G1 | width G1 cycle |
|---|---|---|---|---|---|---|---|---|---|
| DN2 | not possible to simulate with the two populations model (no data on precursors) | | | | | | | | |
| DN3A | No var | Y | full (6 points) | **30.0** [ 29.1 31.0 ] | **21.6** [ 20.6 , 22.6 ] | **8.0** [ 7.6 , 8.3 ] | **0.42** [ 0.4 , 0.5 ] | **74.5** [ 73.1 , 75.8 ] | |
| DN3B | Var G1 | N | full (6 points) | **15.5** [ 15.1 15.8 ] | **8.8** [ 8.5 , 9.1 ] | **5.9** [ 5.7 , 6.1 ] | **0.73** [ 0.7 , 0.8 ] | | **8.8** [ 8.0 , 9.6 ] |
| DN4 | Var G1 | Y | full (6 points) | **12.4** [ 11.9 12.9 ] | **6.3** [ 6.0 , 6.6 ] | **5.5** [ 5.3 , 5.8 ] | **0.57** [ 0.5 , 0.6 ] | **36.8** [ 34.9 , 38.8 ] | **3.9** [ 3.2 , 4.5 ] |
| preSel | Var G1 | Y | full (6 points) | **16.8** [ 16.0 17.5 ] | **9.0** [ 8.2 , 9.7 ] | **7.5** [ 6.9 , 8.0 ] | **0.4** [ 0.4 , 0.4 ] | **87.9** [ 87.0 , 88.8 ] | **2.7** [ 2.1 , 3.3 ] |

**Table S1. Identified durations of cycle phases and variation per population.** The average value for the best set of 10 independent parameter estimations are shown, with the best model hypothesis. The DN2 population required the full dataset (6 time points), while the training data from other populations was already sufficient to identify the cell cycle phase durations. Confidence intervals are shown, obtained from 10 independent parameter estimation after bootstrapping the experimental data (see Methods).

### Table S2

| | Unknown parameters, in hours (same boundaries for all populations) | | | | | | |
|---|---|---|---|---|---|---|---|
| | none var. | G1 var. | S var. | G2M var. | all var. | min | max |
| average G1 (µG1) | *fitted* | *fitted* | *fitted* | *fitted* | *fitted* | 1 | 24 |
| average S (µS) | *fitted* | *fitted* | *fitted* | *fitted* | *fitted* | 1 | 20 |
| average G2M (µG2M) | *fitted* | *fitted* | *fitted* | *fitted* | *fitted* | 0.2 | 5 |
| width G1 (σG1) | 0 | *fitted* | 0 | 0 | *fitted* | 0.1 | 10 |
| width S (σS) | 0 | 0 | *fitted* | 0 | *fitted* | 0.1 | 10 |
| width G2M (σG2M) | 0 | 0 | 0 | *fitted* | *fitted* | 0.1 | 2.5 |
| fraction qiescent (q0) | *fitted in the conditions with long G1* | | | | | 0.01 | 0.98 |

| | Fixed parameters |
|---|---|
| n. of divisions (Ndiv) | 5 |
| death rate (δ, frac. / hour) | 0.0001 |
| population size (NCells) | 10000 |
| threshold EdU+ ∈ [0,2] | 0.01 |
| threshold BrdU+ ∈ [0,2] | 0.01 |
| start EdU pulse (tEdU) | 0 |
| EdU effect duration (θEdU) | 0.75 |
| timeBrdU (tEdU) | 1 |
| durationBrdU (θBrdU) | 0.75 |

**Table S2. Set of parameters defining a dual-pulse simulation for estimating the cell phase durations.** Unknown parameters to be estimated are shown as 'fitted' with their respective minimum and maximum boundaries, depending on the heterogeneity hypothesis and the presence of "long G1" cells. Simulation parameters are also given.

### Table S3

| 1/ decided at birth |
|---|
| time of birth (i.e. start of G1) |
| time of end G1 |
| time of end S |
| time of end G2/M |
| time of death (might happen during the cycle) |
| generation |
| markedForExit = Stay / ExitAfterMitosis |
| **2/ updated during simulation** |
| state(QuiescentG0 / DividingG1 / DividingS / DividingG2M / Dead) |
| current amount of DNA, between 1 (2N) and 2 (4N) |
| current amount of BRDU labeled DNA |
| current amount of EDU labeled DNA |

**Table S3. Description of agent properties used in the simulation.** Events are decided at birth by sampling respective distributions. When daughter cells enter the last division in a population, daughter cells will be marked either for direct exit after completion of the cycle, or for another division, with a probability to have on average Ndiv divisions at the population level. DNA levels are also stored for each agent but are continuously updated during the simulation.

### Table S4

| name | gated on | meaning |
|---|---|---|
| Variables used for training on 4 time-points and validation on the remaining 2 time-points | | |
| **%Mid/late** | | % of EdU+BrdU+ among all cells |
| **%Early** | | % of EdU-BrdU+ among all cells |
| **%Post** | | % of EdU+BrdU- among all cells |
| **%Unstained** | | % of EdU-BrdU- among all cells |
| **Tot in G1** | | % of cells in G1 (irrespective of labelling) |
| **Tot in S** | | % of cells in S (irrespective of labelling) |
| **Tot in G2M** | | % of cells in G2M (irrespective of labelling) |
| **Early in G1** | EdU-BrdU+ | Percent of G1 cells within the early population |
| **Mid/late in G1** | EdU+BrdU+ | Percent of G1 cells within the mid/late population |
| **Post in G1** | EdU+BrdU- | Percent of G1 cells within the Post population |
| **Post in S** | EdU+BrdU- | Percent of S cells within the Post population |
| **Unstained in S** | EdU-BrdU- | Percent of S cells within the Unstained population |
| Variables only used for validation | | |
| Avg DNA Early | EdU-BrdU+ | amount of total DNA (labelled or not) in early cells, rescaled between 0 (2N) and 1 (4N) |
| Avg DNA Mid/Late | EdU+BrdU+ | amount of total DNA (labelled or not) in mid/late cells, rescaled between 0 (2N) and 1 (4N) |

**Table S4: Description of the experimentally observed variables used for estimating the cycle phases that are directly compared to simulation.** Mid/late cells were in the S phase during both labeling periods, while early cells were not yet in the S phase during the first labeling (EdU) and entered the S phase during the second labeling (BrdU). Post cells were in the S phase at first labeling and left before the second labeling. Unstained cells were never in the S phase during the two labeling periods.

### Algorithms 1-4

**Algorithms 1-4:** Algorithmic description of the agent-based model.
**Algorithm 1:** Generation of a new cell at G0, G1 or at a random phase of the cycle.

**Algorithm 1** Generating a new cycling cell with new predefined fate according to the time-distributions, either at a random cycle phase, or in the beginning of its G1.

```
procedure GENERATECELL(t, state = random / G1birth / G0quiescent, generation, DistribG1, DistribS, DistribG2M, DistribDeath)
    A ← new cell (agent)
    A->tbirth ← t
    A->tdie ← A->tbirth + DistribDeath->randValue()
    A->tendG1← A->tbirth + DistribG1->randValue()
    A->tendS← A->tendG1 + DistribS->randValue()
    A->tendG2M← A->tendS + DistribG2M->randValue()
    A->gen ← generation
    A->markForExit = Stay                      ▷ Possible values: Stay / ExitAfterMitosis
    A->cycleState ← DividingG1

    if state == random then          ▷ To start at another phase than G0, randomly shift each phase
        lifespan ← min(tendG2M, tdie)
        shiftFactor ← uniformRandom(0,lifespan)
        A->cycleState ← where 'shiftFactor' lies
        A->tbirth −= shiftFactor
        A->tdie −= shiftFactor
        A->tendG1 −= shiftFactor
        A->tendS −= shiftFactor
        A->tendG2M −= shiftFactor
        A->tdisappear −= shiftFactor
    end if

    if state == G0quiescent then                              ▷ Will stay forever in G0
        A->tdie ← +inf
        A->tendG1← +inf
        A->tendS← +inf
        A->tendG2M← +inf
        A->cycleState ← QuiescentG0
    end if

    A->DNA ← UpdateDNA(time, A->tendG1, A->tendS)    ▷ 1 before S, 2 after S, linear inside S
    A->EdU ← 0                                           ▷ EdU labelled DNA ∈ [0..2]
    A->BrdU ← 0                                          ▷ BrdU labelled DNA ∈ [0..2]
    return A
end procedure

procedure UPDATEDNA(time, tendG1, tendS)
    Return min(2, 1 + max(0, (time−tendG1)/(tendS−tendG1)))
end procedure
```

### Algorithms 1-4

**Algorithms 1-4:** Algorithmic description of the agent-based model.
**Algorithm 2:** Creation of an initial population of cells already at steady-state generations and cycle phases.

**Algorithm 2** Generating a population of cells at equilibrium, spread across generations and cell cycle phases according to the time-distributions, and with a fraction of bystander quiescent cells in G0.

```
procedure INITIALPOPULATION(Ncells, fracQuiescent, Ndiv, DistribG1, DistribS, DistribG2M, DistribDeath, time)
    pop[] <- Empty array of cells                          ▷ Population that will be generated

    fracPerGen ⟵ equilibriumGenerations(Ndiv, DistribG1, DistribS, DistribG2M, DistribDeath)
    n ⟵ ⌊Ndiv⌋                  ▷ Number of 'full' divisions. Generations will go from 0 to n
    for i from 0 to n do
        cumulatedFreqGen[i] ⟵ Ncells (∑_{k=0}^{i} fracPerGen[k]) / (∑_{k=0}^{n} fracPerGen[k]))
    end for

    Nquiescent = ⌊fracQuiescent/Ncells⌋
    for With probability (fracQuiescent / Ncells - Nquiescent) do
        Nquiescent ⟵ Nquiescent + 1                               ▷ 'Smoothing' Nquiescent
    end for

    for nQuiescent times do
        NewCell ⟵ GenerateCell(time, phase=G0quiescent, generation=0)
        pop.append(NewCell)
    end for

    for (Ncell-Nquiescent) times do
        U ⟵ random::uniform(0, 1)           ▷ Will pick a generation according to fracPerGen
        gen ⟵ 0;
        while (gen < n) and (U > cumulatedFreqGen[gen])
            gen ⟵ gen + 1
        end while

        NewCell ⟵ GenerateCell(time, phase=random, generation=gen)
        if gen == n then              ▷ The cells at generation n should not make another cycle
            NewCell->MarkForExit ⟵ exitAfterMitosis
        end if
        pop.append(newCell)
    end for
    Return(pop)
end procedure

Calculates the fraction of cells at each generation when the population reaches equilibium
procedure EQUILIBRIUMGENERATIONS(Ndiv, DistribG1, DistribS, DistribG2M, DistribDeath)
    n ⟵ ⌊Ndiv⌋                                                ▷ Number of 'full' divisions
    T ⟵ DistribG1.mean() + DistribS.mean() + DistribG2M.mean()     ▷ Average cycle duration
    δ = 1/max(10^-10, DistribDeath.mean())          ▷ Death rate, 1/mean of exponential law
    X ⟵ 2(1 − Tδ)                                   ▷ Coefficient of expansion per division
    fractionPerGen ⟵ []
    for i from 0 to n − 1 do
        fractionPerGen[i] ⟵ X^i
    end for
    fractionPerGen[n] ⟵ (Ndiv − n)X^n       ▷ Fraction of cells that complete the nth division
    Return(fractionPerGen)
end procedure
```

### Algorithms 1-4

**Algorithms 1-4:** Algorithmic description of the agent-based model.
**Algorithm 3:** Step-by-step update of the population and cell labeling.

**Algorithm 3** Time-step for evolving the population of cycling cells, and simulating EdU and BrdU labelling according to current levels of EdU and BrdU, and the time distributions

```
procedure TIMESTEP(pop, time, dt, Ncells, Ndiv, EdULevel, BrdULevel, DistribG1, DistribS, DistribG2M, fracQuiescent, DistribDeath)
    for Each cell t in pop do
        if | t->tdie - time | < dt/2 then                                    ▷ Test for death
            t->state ⟵ Dead
            pop.remove(t)
        else
            t->DNA ⟵ updateDNA(time, t->tendG1, t->tendS)
            if t->state == DividingS then                     ▷ Labelling of cells in the S phase
                t -> EdU ⟵ t->EdU + dt.EdULevel / (t->tendS - t->tendG1)
                t -> BrdU ⟵ t->BrdU + dt.BrdULevel / (t->tendS - t->tendG1)
            end if

            if | t->tendG1 - time| < dt/2 then                        ▷ DividingG1 to DividingS
                t->state ⟵ DividingS
            end if
            if | t->tendS - time | < dt/2 then                      ▷ DividingS to DividingG2M
                t->state ⟵ DividingG2M
            end if

            if | t->tendG2M - time | < dt/2 then               ▷ Mitosis to two new cells in G1
                newGen ⟵ t->gen + 1

                for Perform 2 times do                           ▷ Generate two daughter cells
                    Daughter ⟵ generateCell(time, phase=G1birth, newGen)
                    Daughter->EdU ⟵ t->EdU / 2
                    Daughter->BrdU ⟵ t->BrdU / 2
                    if t->markedForExit == stayTilMitosis then       ▷ Parent was marked for exit
                        delete Daughter
                    else
                        if Daughter->gen < Ndiv then        ▷ Daughters at early generations, stay
                            Daughter->markedForExit ⟵ stay
                            pop.append(Daughter)
                        else▷ At last division, daughters stay with probability the decimal part of Ndiv
                            if With probability Ndiv − ⌊Ndiv⌋, do then
                                Daughter->markedForExit ⟵ stayTilMitosis  ▷ Stay until next mitosis
                                pop.append(Daughter)
                            else              ▷ Will exit now and not perform one more cell cycle
                                delete Daughter
                            end if
                        end if
                    end if
                end for                                               ▷ Two daughters generated
                pop.remove(t)             ▷ Remove the parent cell (replace by the two daughters
            end if
        end if
    end for

    Adding constant inflow needed to maintain population at equilibrium:
    T ⟵ DistribG1.mean() + DistribS.mean() + DistribG2M.mean()      ▷ Average cycle duration
    nGen0 ⟵ Ncells . equilibriumGenerations(Ndiv, DistribG1,...)[0]
    inflow ⟵ dt . (1 - fracQuiescent) . nGen0/T      ▷ Inflow to maintain cells amount at gen = 0
    for Repeat inflow times + once with probability (inflow - ⌊inflow⌋) do
        newCell ⟵ generateCell(time, phase=random, gen=0)
        pop.insert(newCell)
    end for
end procedure
```

### Algorithms 1-4

**Algorithms 1-4:** Algorithmic description of the agent-based model.
**Algorithm 4:** Main organization of a simulation returning the cost of a parameter set.

**Algorithm 4** Simulation of a full EdU-BrdU dual pulse experiment

```
procedure OneSimulation(Ncells, fracQuiescent, dt, timeEnd, DistribG1, DistribS, DistribG2M, DistribDeath, EdUdynamics, BrdUdynamics, datasets, thresholdEdU, thresholdBrdU)
    pop ⟵ initializePopulation(Ncells, fracQuiescent, Ndiv, time=0)
    cost ⟵ 0
    for time from 0 to timeEnd by steps of dt do
        EdULevel ⟵ EdUdynamics(time)
        BrdULevel ⟵ BrdUdynamics(time)
        exitedCells ⟵ timeStep(pop, time, dt, Ncells, EdULevel, BrdULevel, DistribG1, ...)
        cost ⟵ cost + compareReadouts(pop, time, datasets, thresholdEdU, thresholdBrdU)
    end for
    Return(cost)
end procedure

procedure compareReadouts(pop, time, datasets, thresholdEdU, thresholdBrdU)
    cost ⟵ 0
    for Each cell t in pop do
        EdUpositive ⟵ t->EdU > thresholdEdU
        BrdUpositive ⟵ t->BrdU > thresholdBrdU

        Calculation/Update of statistics for each type of observable
        Fraction of cells EdU+, BrdU+, and Edu+/-Brdu+/- among pop
        Percent of EDU+/-Brdu+/- population, that are at G0 or G1 phase
        Average DNA levels among EDU+/-Brdu+/- populations
        Fraction of cells in each phase of the cycle
    end for
    if dataset contains datapoints at t=time then
        cost ⟵ cost + statisticalComparison(dataset(time), calculated statistcs)
    end if
    Return(cost)
end procedure
```
